# Sequence termination cues drive habits via dopamine-mediated credit assignment

**DOI:** 10.1101/2024.10.16.618735

**Authors:** Robin Magnard, Yifeng Cheng, Joanna Zhou, Haley Province, Nathalie Thiriet, Patricia H. Janak, Youna Vandaele

**Author notes:** **CORRESPONDENCE** Dr. Youna Vandaele, Université de Poitiers, INSERM, U-1084, Laboratoire des Neurosciences Expérimentales et Cliniques, Poitiers, France., *Email address*, Dr. Robin Magnard, Department of Psychological and Brain Sciences, Johns Hopkins University, Baltimore, MD, USA., *Email address.

## Abstract

**Background:** Mesolimbic dopamine (DA) neurons are central to cue guided reward seeking and action sequence learning. Yet, the mechanisms by which cue-induced DA neural activity drives goal-directed or habitual sequence execution remain unknown.

**Methods:** We designed two novel tasks to isolate the effect of sequence-delineating cues on DA-driven behavioral strategies and learning. In the lever insertion fixed-ratio 5 task (LI5), the lever insertion marked sequence initiation. In the lever retraction fixed-ratio 5 task (LR5), the lever retraction served as both sequence termination and reward-predictive cue.

**Results:** We found that sequence initiation and termination cues differentially affect reward expectation during action sequences, with only the termination cue contributing to greater outcome devaluation insensitivity, automaticity and behavioral chunking. Mesolimbic fiber photometry recording revealed that this habit-like behavior was associated with a rapid backpropagation in DA signals from the reward to the immediately preceding cue and with attenuated DA reward prediction error signals, which reflected greater behavioral inflexibility. Finally, in absence of external cues, brief optogenetic stimulation of VTA DA neurons at sequence termination was sufficient to drive automaticity and, to some extent, behavioral chunking.

**Conclusion:** Our results highlight the critical role of cue-evoked DA signals at sequence termination in mediating credit assignment and driving the development of habitual action sequence execution.

## INTRODUCTION

Mesolimbic dopamine (DA) neurons play a critical role in cue-mediated reward expectation and sequence learning^1–5^. Notably, midbrain DA neurons encode reward prediction errors (RPE)^1–8^, which are deviations from expected outcome that drive learning by conferring motivational value to actions and reward-predictive cues^3,8^. Across sequence learning, DA signals in the nucleus accumbens (NAc) backpropagate from the reward to sequence initiation^9–12^. Mesolimbic DA activity predicts action sequences performance^9,10^ and scale with distance to reward to sustain motivation toward goals^13,14^. Although DA has been related to habit learning and response automaticity^15–19^, it remains unclear how DA responses at action and reward-predictive cues evolve over sequence learning when behavior is goal-directed, i.e. driven by outcome representation, and when it becomes automated and habitual. Indeed, studies of DA dynamics during sequence learning have not characterized whether behavior was habitual or goal-directed ^9–12^, with one exception ^20^, which lacked within-session resolution during early habit formation.

Habits are commonly defined as stimulus-response behaviors^21^, and are predominantly identified as an absence of goal-directed control (i.e. insensitivity to outcome devaluation or contingency degradation)^22,23^. This description has been extended to include chunked action sequences executed automatically without outcome expectation until completion^24–26^. Several studies reveal the importance of sequence-delineating cues in habit learning and behavioral chunking^27–32^, either by making stimulus-response associations more explicit and/or by reducing attentional demand during execution of behavior. We previously showed that in a discrete-trials task, lever insertion, which signals trial onset and sequence initiation, and retraction, which signals reward delivery and sequence termination, promote habit, automaticity and behavioral chunking^27,28,33^. Yet, it remains unclear whether these cues differentially shape the development of these behaviors. It is also unknown whether their presentation differentially influence reward expectation and is associated with different distinct mesolimbic DA activity patterns reflecting specific response strategies.

To assess how sequence initiation and termination cues influence habitual or goal-directed response strategy, we designed two isomorphic fixed-ratio-5 tasks in which only one sequence-delineating cue was present, either lever insertion in LI5 task, or lever retraction in LR5 task. Based on prior work^28^, we hypothesized that the sequence termination cue in the LR5 task would serve as response feedback, alleviating requirements for reward monitoring, thereby promoting automatic and chunked responses. We further predicted that this proximal reward-predictive cue would drive core features of habits such as insensitivity to both outcome devaluation and reversed instrumental contingency^27,28,33^.

We propose that DA-mediated credit assignment^34–36^ may link the sequence termination cue to habit formation by shifting the expectation of reinforcement from individual presses to the entire sequence as a chunk executed automatically. To test this, we tracked mesolimbic DA neuron activity along the action sequence in the LI5 and LR5 task, and across training. Finally, we used optogenetic manipulation to demonstrate the role of DA-mediated credit assignment in the development of automaticity and behavioral chunking.

## MATERIALS & METHODS

Wild-type (WT) (n = 48) or TH-Cre Long-Evans rats (n = 36) were used in this study (details in supplement methods). We designed two fixed-ratio 5 tasks to discern the respective role of initiation (lever insertion) and termination (lever retraction) cues in action sequence learning. In the lever insertion fixed-ratio 5 (LI5) task, trials began with lever insertion. Rats had to complete a sequence of five lever presses to obtain the reward (0.1 ml 20% sucrose), with the lever retracting randomly during reward retrieval and thus not predicting sequence completion (**Fig. 1a**). In contrast, in the lever retraction fixed-ratio 5 (LR5) task, the lever retracted immediately after the fifth lever press, serving as the termination cue to signal reward delivery, while the lever insertion occurred during reward retrieval and did not signal trial onset (**Fig. 1b**).

**Figure 1.**
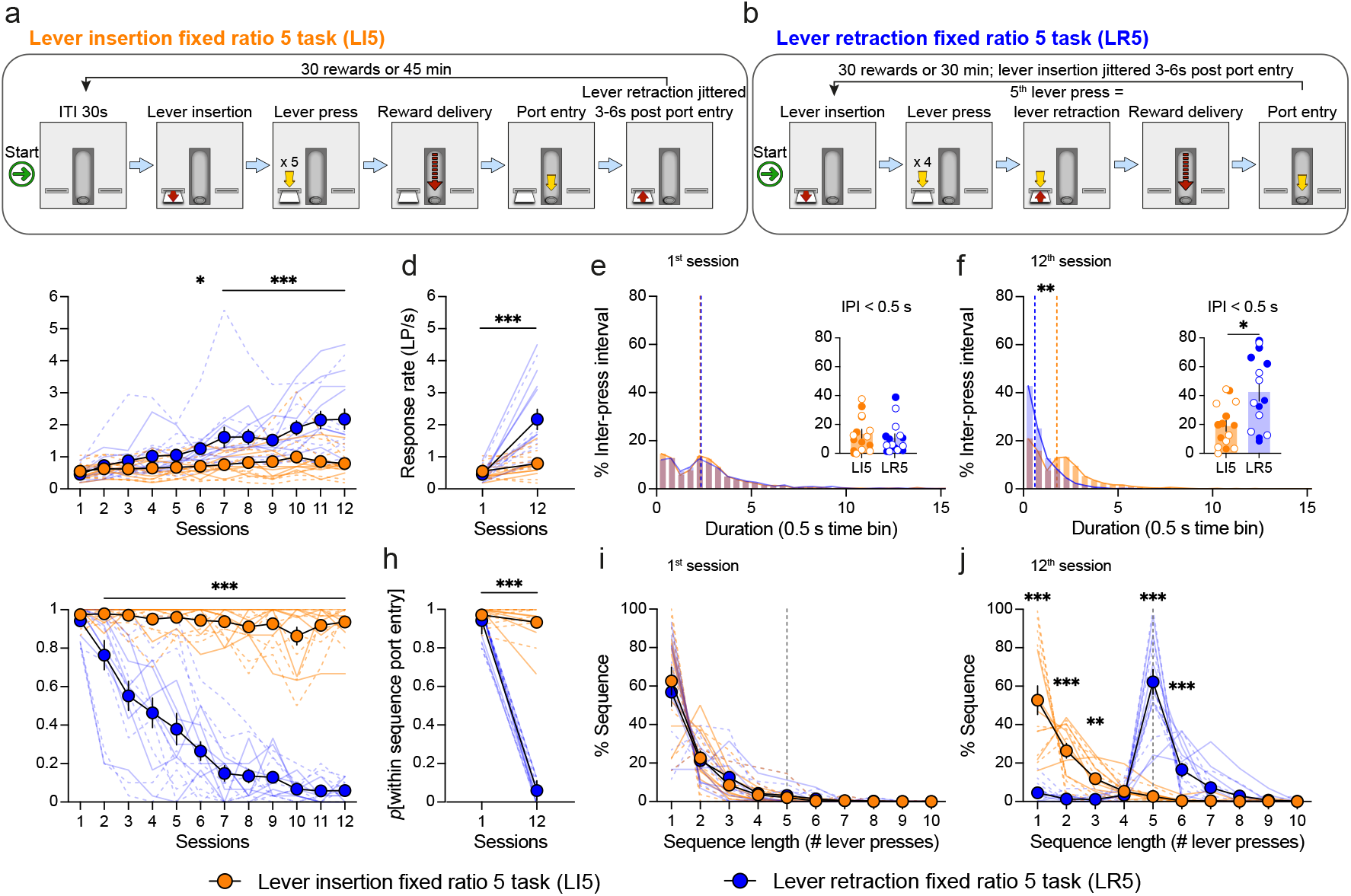
The lever retraction cue, but not the lever insertion cue, promotes action automaticity and behavioral chunking. (**a**) Lever insertion fixed-ratio 5 (LI5) task design. (**b**) Lever retraction fixed-ratio 5 (LR5) task design. (**c**) Response rate in lever press per second (LP/s), within the lever press sequence during LI5/LR5 training sessions (two-way RM ANOVA, task effect: F_1,28_ = 10.6, p = 0.003; task x session interaction: F_11,308_ = 10.07, p < 0.001; Student Newman-Keuls post hoc pairwise comparison). (**d**) Response rate on the first versus last session in the LI5/LR5 tasks (two-way RM ANOVA, task effect: F_1,28_ = 11.54, p = 0.002; task x session interaction: F_1,28_ = 23.88, p < 0.001; Student Newman-Keuls post hoc pairwise compari-son). (**e**) Frequency distribution of inter-press intervals (IPI) for the 1st LI5/LR5 session (Mann-Whitney Rank Sum t-test median comparison U = 455, p = 0.725). Insert panel: individual proportion of IPI under 0.5 s (Mann-Whitney Rank Sum t-test, U = 94, p = 0.455). (**f**) Frequency distribution of IPI for the 12th LI5/LR5 session (Mann-Whitney Rank Sum t-test median comparison U = 290, p = 0.007). Insert panel: individual proportion of IPI under 0.5 s (Mann-Whitney Rank Sum t-test, U = 51, p = 0.011). (**g**) Probability to enter the port within the lever press sequence (within-sequence port entry) during training in LI5/LR5 tasks (two-way RM ANOVA, task effect: F_1,28_ = 247.71, p < 0.001; task x session interaction: F_11,308_ = 39.95, p < 0.001; Student Newman-Keuls post-hoc pairwise comparison). (**h**) Probability of within sequence port entry on the first versus last training session in the LI5/LR5 tasks (two-way RM ANOVA, task effect: F_1,28_ = 483.56, p < 0.001; task x session interaction: F_1,28_ = 687.4, p < 0.001; Student Newman-Keuls post hoc pairwise comparison). (**i**) Frequency distribution of sequence lengths during the first training session in the LI5/LR5 tasks. The grey dashed line represents the optimal sequence length of 5 lever presses (two-way RM ANOVA, no task effect: F_1,28_ < 1, p = 0.808; no task x sequence length interaction: F_9,252_ < 1, p = 0.95). (**j**) Frequen-cy distribution of sequence length during the last training session in the LI5/LR5 tasks. The grey dashed line represents the optimal sequence length of 5 lever presses (three-way RM ANOVA, no task effect: F_1,28_ < 1, p = 0.919; task x sequence length interaction: F_9,252_ = 49.1, p < 0.001; Student Newman-Keuls post-hoc pairwise comparison). Data shown as mean ± SEM, supe-rimposed with individual data point. LI5: n = 15 (7 males; 8 females); LR5: n = 15 (7 males; 8 females). Solid lines and filled circles: males; Dashed lines and empty circles: females. ITI: inter-trial interval. *p < 0.05; **p < 0.01; ***p < 0.001.

The first experiment aimed at characterizing behavior in the two tasks. To measure habit learning, sensitivity to outcome devaluation was assessed after 12 training sessions. In the second and third experiments, we used fiber photometry to respectively record ventral tegmental area (VTA) DA neural activity and DA release in the NAc Core. Habit learning was assessed by testing response sensitivity to an omission schedule. DA signals were recorded throughout training and during the omission test. In the fourth experiment, TH-Cre^+^ rats received optogenetic VTA DA neurons stimulation at sequence completion and in absence of lever retraction cue for 12 consecutive sessions. Virus expression and optic fiber placement were verified with histological procedures. For full methodological details, see supplement methods.

## RESULTS

### Sequence termination cues promote habit-like behavior

Task differences rapidly emerged during training with increased response rate for rats trained in the LR5 task, compared to the LI5 task (**Fig. 1c-d**). While both groups had identical inter-press intervals (IPI) in the first session, with a similar proportion of IPIs shorter than 0.5s (**Fig. 1e**), LR5 rats showed a greater proportion of shorter IPIs on the last training session (**Fig. 1f**), indicating greater action automaticity. Consistent with these observations, LR5 rats rapidly learned to continuously press on the lever until its retraction, leading to a drastic decrease in the likelihood of port checking during sequence execution (**Fig. 1g**). In contrast, LI5 rats maintained a high probability of port-checking across sessions (**Fig. 1g-h**). This pattern suggests that LR5 rats learned to concatenate individual lever presses into a single behavioral sequence executed without intermediate reward expectation, a hallmark of behavioral chunking. Early in training, LR5 rats often check the port after 1-3 lever presses (**Fig. 1i**), but by training end, they reliably completed full lever press sequences without checking (**Fig. 1j**). In contrast, LI5 rats showed no evidence of chunking, as they persisted in port checking after each response throughout training (**Fig. 1i** and **1j**).

Task differences in action automaticity and behavioral chunking were associated with differences in sensitivity to sensory-specific satiety-induced outcome devaluation (**Suppl. Fig. 1**). Indeed, we observed a higher proportion of rats insensitive to outcome devaluation in the LR5 task compared to the LI5 task (devaluation ratio ≤ 0; 8/15 in LR5 task vs 1/15 in LI5 task). We found no effect of sex in these tests (**Suppl. table 1**). Taken together, these results show that the lever retraction cue, rather than the lever insertion cue, promotes habit-like behavior, characterized by action automaticity, behavioral chunking, and a reduced sensitivity to outcome devaluation.

### Distinct patterns of VTA DA neuron activity in response to cues and reward depending on response strategy

To determine if DA dynamics during sequence execution depend on cue-mediated reward expectation and prediction, we used fiber photometry to record VTA DA neuron activity across training in the LR5 and LI5 tasks. GCaMP6f was expressed in (TH)-Cre^+^ rats to selectively target VTA DA neurons with 95% specificity (**Fig. 2a-c**). We replicated the task differences in behavior (**Suppl. Fig. 2**). We observed no sex differences in behavior or neural activity and combined both sexes in the following analyses (**Suppl. table 1**).

**Figure 2.**
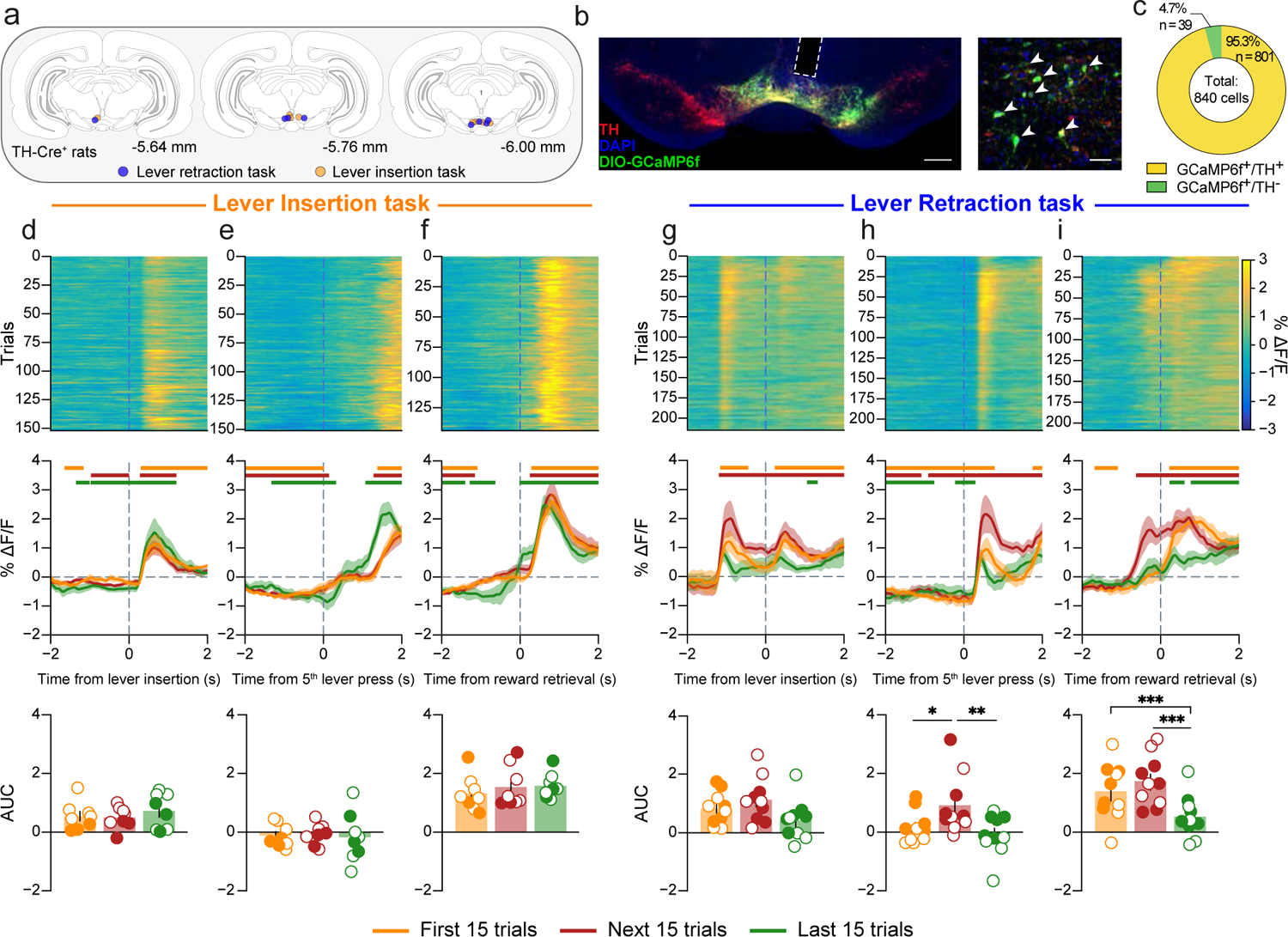
Training-dependent changes in VTA dopamine neuronal phasic activity in response to cues and reward in the LR5 task, but not the LI5 task. (**a**) Coronal sections showing location of optic fiber tips relative to Bregma for TH-Cre+ rats in LI5 and LR5 tasks. (**b**) Left panel: DIO-GCaMP6f was infused in the VTA of TH-Cre^+^ rats, and the optic fiber, represented with the white dash line, was implanted above the infusion site. Scale bar: 500 µm. Right panel: histology confirmation of DIO-GCaMP6f (green) and TH (red) co-expression (indicated by white arrows). Scale bar: 50 µm. (**c**) GCaMP6f virus is predo-minantly expressed in TH^+^ neurons. (**d**) Top panel: LI5 task heatmap displaying trial-by-trial dopamine neuron activity, averaged across rats, around lever insertion. Middle panel: phasic dopamine neuron activity around lever insertion averaged across rats and trials for the first 15 trials (orange line), the next 15 trials (red line) and the final 15 trials (green line) of training. Bottom panel: area under the curves (AUCs) for 0 to 1 s following lever insertion (one-way RM ANOVA, no trial effect: F_2,14_ < 1, p = 0.43). (**e**) Same as (**d**) for the 5th lever press event in the LI5 task (one-way RM ANOVA, no trial effect: F_2,14_ < 1, p = 0.863). (**f**) Same as (**d**) for reward retrieval in the LI5 task (one-way RM ANOVA, no trial effect: F_2,14_ = 2.29, p = 0.147). (**g**) Same as (**d**) for lever insertion in the LR5 task (one-way RM ANOVA, no trial effect: F_2,18_ = 2.71, p = 0.09). (**h**) Same as (**d**) for the 5th lever press in the LR5 task (one-way RM ANOVA, trial effect: F_2,18_ = 7.26, p = 0.005; Student Newman-Keuls post hoc pairwise comparison). (**i**) Same as (**d**) for reward retrieval in the LR5 task (one-way RM ANOVA, trial effect: F_2,18_ = 21.04, p < 0.001; Student Newman-Keuls post hoc pairwise comparison). Data shown as means ± SEM, superimposed with individual data point. LI5 n = 8 (3 males; 5 females); LR5 n = 10 (5 males; 5 females). Orange, red and green horizontal bars indicate significant deviation from baseline at 99% confidence level. Filled circles: males; Empty circles: females. *p < 0.05; **p < 0.01; ***p < 0.001.

We monitored DA neural activity at lever insertion, lever retraction, during lever presses, and at reward retrieval. We analyzed changes in activity across 3 sets of 15 trials: the first 15 trials of the first session (trials 0-15), the second 15 trials of the first session (trials 16-30), and the last 15 trials of the last training session.

In the LI5 task, where lever insertion signals opportunity to respond for reward, DA neural activity increased at lever insertion (**Fig. 2d**) and at reward retrieval (**Fig. 2f**), with no change across trials and no response at sequence completion (**Fig. 2e**). In contrast, in the LR5 task, where lever retraction signals sequence completion and reward delivery, we observed rapid training-related changes in DA activity shifting from reward to the retraction cue as learning progressed (**Fig. 2h-i**), and no training-related changes at lever insertion (**Fig. 2g**). With extended training in the LR5 task, both the lever retraction and reward-related DA responses declined (**Fig. 2h-i**; last 15 trials).

### Distinct patterns of VTA DA neuron activity during lever press sequences between the LI5 and LR5 tasks

DA neural activity patterns during the lever press sequence differed across tasks (**Fig. 3**). To compare activity at lever presses, we examined the first 4 responses as the fifth press coincides with lever retraction in the LR5 task (**Fig. 2**). In the LI5 task, DA neural activity associated with sequence initiation changed over training, initially peaking after the first lever press (first and next 15 trials) and progressively shifting to before the lever press (last 15 trials; **Fig. 3a**). DA response decreased as rats progressed in the sequence from the first to fourth response (**Fig. 3e-f**), with an overall suppression immediately post-press during the last 15 trials (**Fig. 3 a-d and g**). Similarly, phasic DA responses evoked by premature port-checking also declined along the sequence (**Suppl. Fig. 3a-d**). Notably, 1-2s after each port entry, we observed an increased inhibition in DA neural activity along the sequence (**Suppl. Fig. 3e-g**). This inhibition in LI5-trained rats might reflect a negative reward prediction error (RPE) signal, indicating a growing violation of reward expectation as rats approached sequence completion.

**Figure 3.**
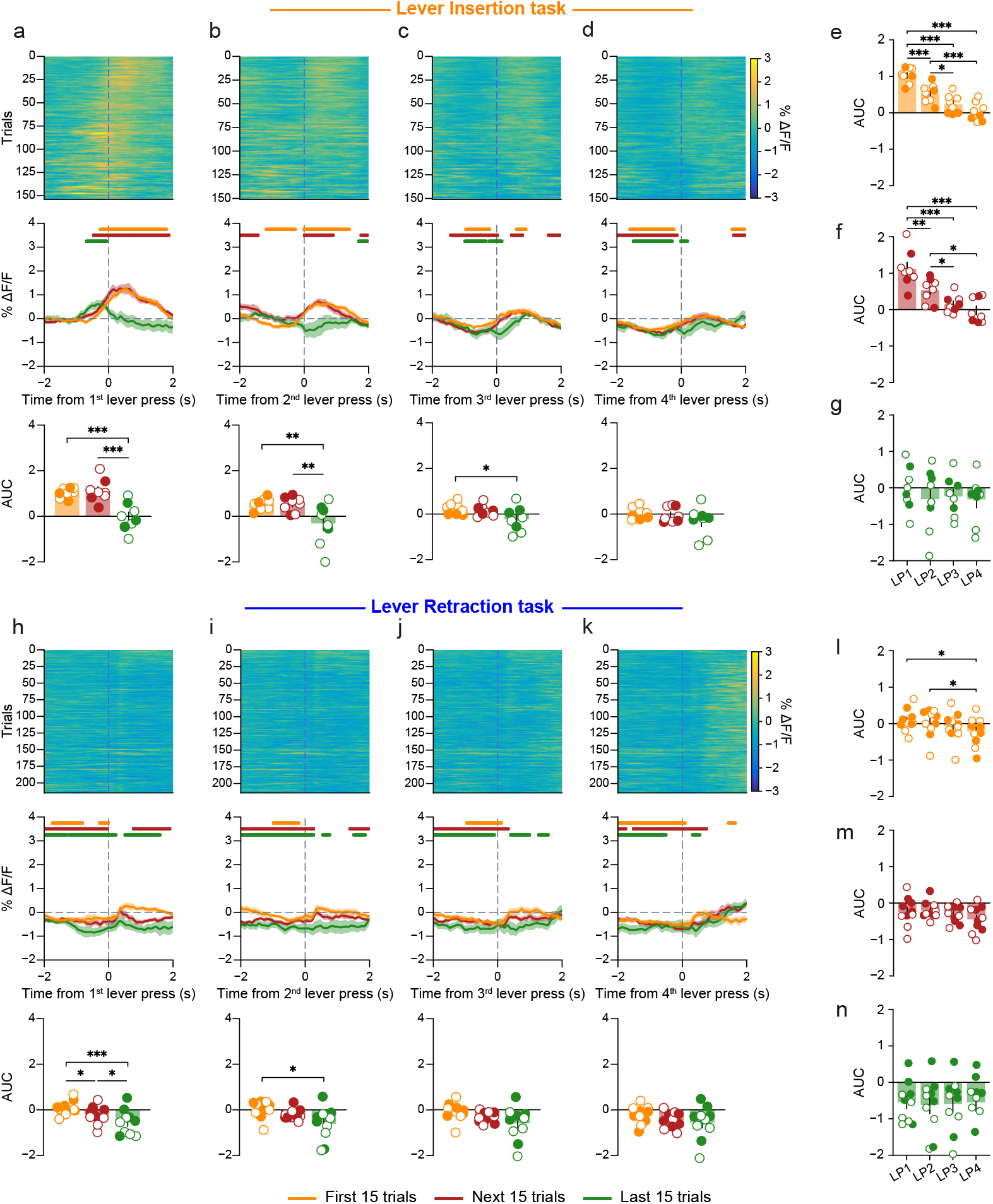
Distinct patterns of VTA DA neuronal activity during lever press sequences in the LI5 and LR5 tasks. (**a-d**) Sequence of the first four lever presses for the LI5 task. The upper panel heatmaps illustrate trial-by-trial dopamine neuron activity, averaged across rat. The middle panel represents dopamine neuron activity averaged across rats and trials, delineated for the first 15 trials (orange line), the next 15 trials (red line), and the final 15 trials (green line) of the training period. The bottom panel displays the area under the curves (AUCs) for the 0 to 1 seconds following the event. (**a**) 1st lever press (LP1; one-way RM ANOVA, trial effect: F_2,14_ = 14.71, p < 0.001. Student Newman-Keuls post hoc pairwise comparison). (**b**) 2nd lever press (LP2; one-way RM ANOVA, trial effect: F_2,14_ = 9.55, p = 0.002. Student Newman-Keuls post hoc pairwise comparison). (**c**) 3rd lever press (LP3; one-way RM ANOVA, trial effect: F_2,14_ = 4.54, p = 0.03. Student Newman-Keuls post hoc pairwise comparison). (**d**) 4th lever press (LP4; one-way RM ANOVA, no trial effect: F_2,14_ = 2.09, p = 0.160). (**e-g**) Area under the curves (AUCs) for the 0 to 1 s window following the first four lever presses of the sequence in the LI5 task, measured across the three trial periods (two-way RM ANOVA, trial effect: F_2,14_ =10.12, p = 0.002; lever press effect: F_3,21_ = 28.63, p < 0.001; lever press x trial interaction F_6,42_ = 5.77, p < 0.001). (**e**) First 15 trials (one-way RM ANOVA, lever press effect: F_3,21_ = 33.05, p < 0.001. Student Newman-Keuls post hoc pairwise comparison). (**f**) Next 15 trials (one-way RM ANOVA, lever press effect: F_3,21_ = 17.71, p < 0.001. Student Newman-Keuls post hoc pairwise comparison). (**g**) Final 15 trials (one-way RM ANOVA, lever press effect: F_3,21_ = 2.22, p = 0.116). (**h-k**) same as (**a-d**) for the LR5 task. (**h**) 1st lever press (LP1; one-way RM ANOVA, trial effect: F_2,18_ = 9.95, p < 0.001. Student Newman-Keuls post hoc pairwise comparison). (**i**) 2nd lever press (LP2; one-way RM ANOVA, trial effect: F_2,18_ = 4.60, p = 0.024. Student Newman-Keuls post hoc pairwise comparison). (**j**) 3rd lever press (LP3; one-way RM ANOVA, no trial effect: F_2,18_ = 2.95, p = 0.08). (**k**) 4th lever press (LP4; one-way RM ANOVA, no trial effect: F_2,18_ = 1.57, p = 0.234). (**l-n**) same as (**e-g**) for the LR5 task (two-way RM ANOVA, trial effect: F_2,18_ = 4.77, p = 0.02; lever press effect: F_3,27_ = 3.23, p = 0.04; no lever press x trial interaction F_6,54_ = 1.25, p = 0.29). (**l**) First 15 trials (one-way RM ANOVA, lever press effect: F_3,27_ = 4.12, p = 0.015. Student Newman-Keuls post hoc pairwise comparison). (**m**) Next 15 trials (one-way RM ANOVA, lever press effect: F_3,27_ = 1.55, p = 0.225). (**n**) Final 15 trials (one-way RM ANOVA, lever press effect: F_3,27_ < 1, p = 0.824). Data shown as means ± SEM, superimposed with individual data point. LI5 n = 8 (3 males; 5 females); LR5 n = 10 (5 males; 5 females). Orange, red and green horizontal bars indicate significant deviation from baseline at 99% confidence level. Filled circles: males; Empty circles: females. *p < 0.05; **p <0.01; ***p < 0.001.

In contrast, LR5-trained rats showed minimal DA modulations around lever presses (**Fig. 3h-k**), with modest decreases across training (**Fig. 3l-n**). Deconvolution analysis, isolating neural responses to specific events while avoiding contamination from adjacent events (supplement methods), confirmed similar patterns of DA neural activity between the first and last training sessions in both tasks (**Suppl. Fig. 4)**.

### Patterns of DA release in the Nucleus Accumbens Core in response to cues, lever presses and reward are similar to VTA

We next asked if the distinct patterns of VTA DA neuron activity were similarly observed in DA release in the NAc Core, a primary target for VTA DA neurons implicated in DA-mediated cue processing and reward learning^37–40^. To this end, we infused the DA sensor dLight1.2 into NAc Core in male and female rats (**Fig. 4a-b**).

**Figure 4.**
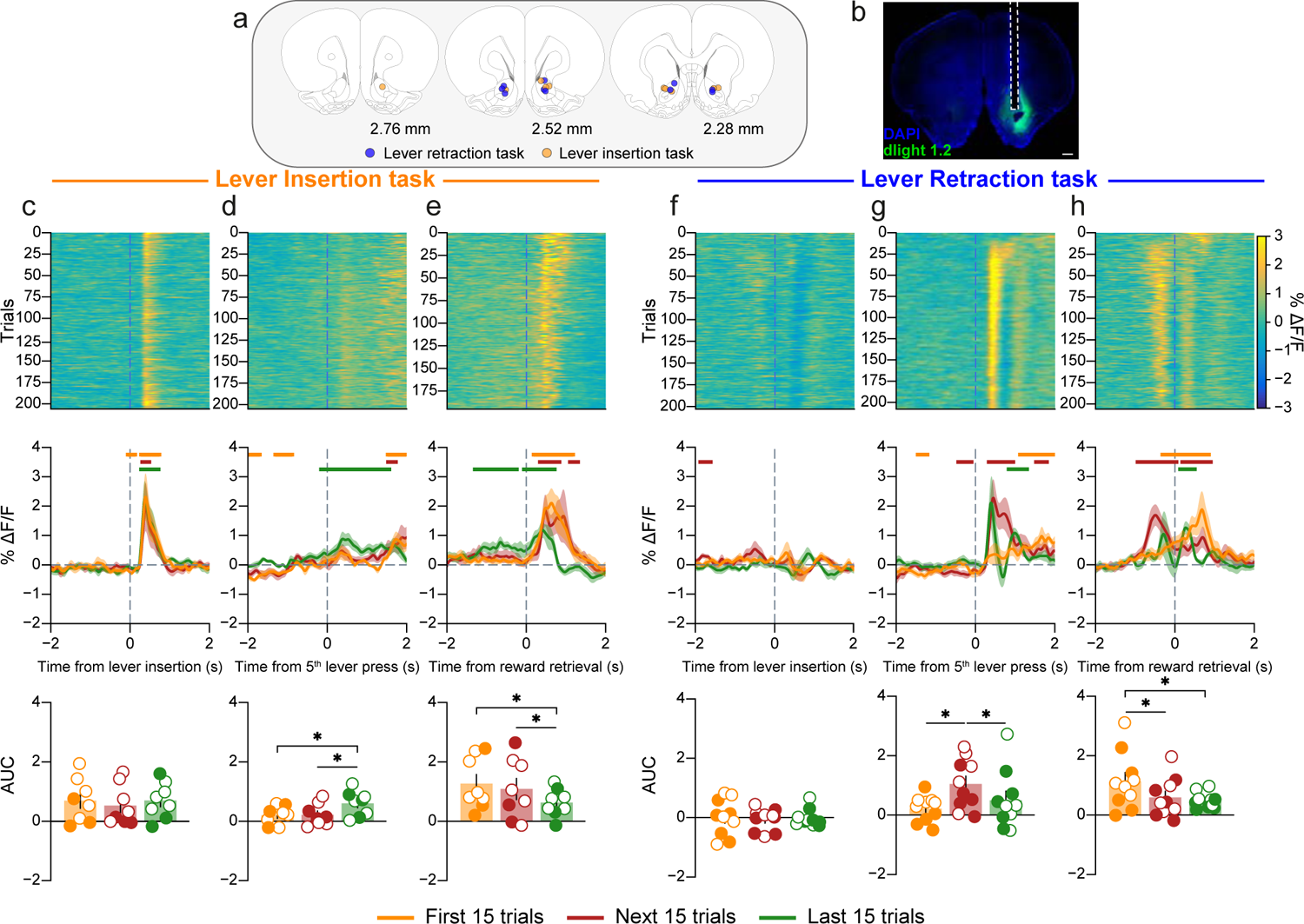
Dynamic changes in dopamine release in the NAc Core in response to cues and reward in the LI5 and LR5 tasks. (**a**) Coronal sections showing location of optic fiber tips relative to Bregma in LI5 and LR5 tasks. (**b**) dLight 1.2 was infused in the NAc Core, and the optic fiber, represented with the white dash line, was implanted above the injection site. Scale bar: 500 µm. (**c**) Top panel: LI5 task heatmap displaying trial-by-trial dopamine release, averaged across rats, around lever insertion. Middle panel: phasic dopamine release around lever insertion averaged across rats and trials for the first 15 trials (orange line), the next 15 trials (red line) and the last 15 trials (green line) of training. Bottom panel: area under the curves (AUCs) for 0 to 1 seconds following lever insertion (one-way RM ANOVA, no trial effect: F_2,14_ < 1, p = 0.59). (**d**) Same as (**c**) for the 5th lever press event in the LI5 task (one-way RM ANOVA, trial effect: F_2,14_ = 5.29, p = 0.02; Student Newman-Keuls post hoc pairwise comparison). (**e**) Same as (**c**) for reward retrieval in the LI5 task (one-way RM ANOVA, trial effect: F_2,14_ = 5.42, p = 0.02; Student Newman-Keuls post hoc pairwise comparison). (**f**) Same as (**c**) for lever insertion in the LR5 task (one-way RM ANOVA, no trial effect: F_2,18_ < 1, p = 0.86). (**g**) Same as (c) for the 5th lever press in the LR5 task (one-way RM ANOVA, trial effect: F_2,18_ = 5.53, p = 0.013; Student Newman-Keuls post hoc pairwise comparison). (**h**) Same as (**c**) for reward retrieval in the LR5 task (one-way RM ANOVA, trial effect: F_2,18_ = 6.23, p = 0.009; Student Newman-Keuls post hoc pairwise comparison). Data shown as means ± SEM, superimposed with individual data point. LI5 n = 8 (3 males; 5 females); LR5 n = 10 (5 males; 5 females). Orange, red and green horizontal bars indicate significant deviation from baseline at 99% confidence level. Filled circles: males; Empty circles: females. *p < 0.05.

DA release patterns tended to mirror VTA DA neuron activity across training. The lever insertion cue elicited a peak in DA release in the LI5 task (**Fig. 4c**) but not in the LR5 task (**Fig. 4f**). Conversely, the fifth lever press did not evoke DA activity in the LI5 task (**Fig. 4d**), while it caused a significant DA release in the LR5 task (**Fig. 4g**). In addition, LI5 rats showed a reward-evoked DA release that somewhat diminished by training end (**Fig. 4e**), whereas in LR5 rats, the reward-related peak rapidly shifted to the lever retraction cue (**Fig. 4h**).

Consistent with VTA recordings, LI5 rats exhibited DA release time-locked to lever presses (**Suppl. Fig. 5a-g**), but the amplitude of the DA signal remained stable across training. No lever press-elicited DA release was observed in LR5-trained rats (**Suppl. Fig 5h-n**).

Combining VTA and NAc Core recordings, we found that in the LR5 task, where lever retraction signals sequence completion, DA responses rapidly shift within a single session, from reward retrieval to sequence termination. In contrast, in the LI5 task, where no cue signals reward delivery, DA signals remain anchored to the reward, even after extended training. Importantly, recurrent port checking in LI5 task likely reflects progress monitoring. Together, these results illustrate how animals adapt distinct behavioral strategies to achieve their goals based on the sequence-delineating cue presented.

### Impaired contingency learning induced by the lever retraction cue is associated with attenuated RPE DA signals in the VTA and NAc Core

After training in the LR5 and LI5 tasks, rats underwent a single session under an omission schedule to evaluate behavioral and dopaminergic responses to reversed instrumental contingencies. The omission procedures mirrored the task structures experienced during training, but required rats to refrain from pressing the lever for 20 seconds to receive a reward (**Fig. 5a-b, supplement methods**).

**Figure 5.**
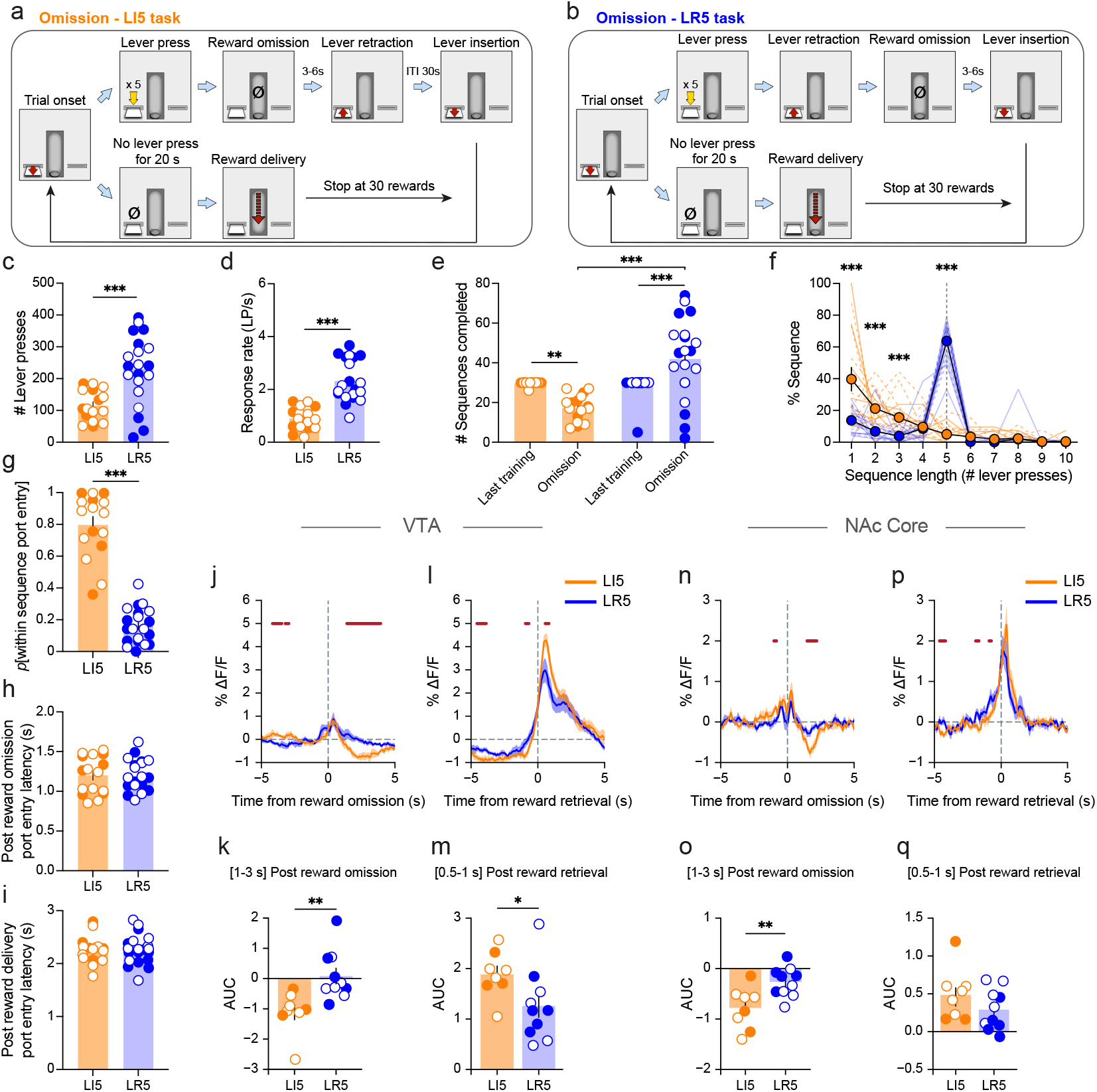
Impaired contingency learning induced by the lever retraction cue is associated with attenuated RPE DA signals in the mesolimbic pathway. Omission test design for the LI5 task (**a**) and the LR5 task (**b**). (**c**) Total number of lever presses during the omission test (two-way ANOVA, task effect: F_1,32_ = 17.87, p < 0.001; no task x region interaction: F_1,32_ < 1, p = 0.529). (**d**) Response rate in lever press per second (LP/s), within the lever press sequence (two-way ANOVA, task effect: F_1,32_ = 40.49, p < 0.001; no task x region interaction: F_1,32_ < 1, p = 0.821). (**e**) Number of sequences completed during the last LI5/LR5 training session and the omission test (three-way RM ANOVA, task effect: F_1,32_ = 18.39, p < 0.001; task x session interaction: F_1,32_ = 34, p < 0.001; no task x session x structure interaction F_1,32_ = 1.43, p = 0.24). (**f**) Frequency distribution of sequence lengths during the omission session. The grey dashed line represents the “optimal” sequence length of 5 lever presses (three-way RM ANOVA, no task effect: F_1,32_ < 1, p = 0.585; task x sequence length interaction: F_9,288_ = 47.06, p < 0.001; no task x sequence length x region interaction: F_9,288_ < 1, p = 0.979; Student Newman-Keuls post hoc pairwise comparison). (**g**) Probability to enter the port within the lever press sequence (within-sequence port entry) (two-way ANOVA, task effect: F_1,32_ = 155.75, p < 0.001; no task x region interaction: F_1,32_ = 1.98, p = 0.169). (**h**) Port entry latency following omission of the reward (two-way ANOVA, no task effect: F_1,32_ < 1, p = 0.95; no region effect: F_1,32_ < 1, p = 0.74; task x region interaction: F_1,32_ = 4.41, p = 0.044). (**i**) Port entry latency following unexpected reward delivery (two-way ANOVA, no task effect: F_1,32_ < 1, p = 0.940; no task x region interaction: F1,32 < 1, p = 0.361). (**j**) Phasic VTA dopamine neuron activity around port entries following reward omission. (**k**) Area under the curves for 1 to 3 seconds after reward omission (unpaired two-tailed t-test comparison, t_16_ = -3.30, p = 0,004). (**l**) Phasic VTA neuron activity around reward retrieval. (**m**) Area under the curves for 0.5 to 1 seconds after reward retrieval (unpaired two-tailed t-test comparison, t16= 2.15, p = 0.047). (**n**) Phasic NAc Core DA signal around port entries following reward omission. (**o**) Area under the curves for 1 to 3 seconds after reward omission (unpaired two-tailed t-test comparison, t16= -3.07, p = 0,007). (**p**) Phasic NAc Core DA signal around reward retrieval. (**q**) Area under the curves for 0.5 to 1 seconds after reward retrieval (unpaired two-tailed t-test comparison, t_16_ = 1.36, p = 0.191). Data shown as means ± SEM, superimposed with individual data point. LI5 n = 16 (6 males; 10 females); LR5 n = 20 (10 males; 10 females). Red horizontal bars indicate significant difference between LI5 and LR5 tasks at 95% confidence level. Filled circles: males; Empty circles: females. *p < 0.05; **p < 0.01; ***p < 0.001.

During the omission test, LR5-trained rats responded more than LI5 rats (**Fig. 5c**), indicating reduced ability to adapt to the new contingencies. LR5 rats completed the 5-lever press sequence faster than LI5 rats (**Fig. 5d**). LI5 rats completed fewer sequences in the omission test compared to the last training session whereas LR5 rats completed more sequences (**Fig. 5e**). While LI5 rats frequently checked the magazine between responses, LR5 rats remained cue-driven and completed most sequences in chunks of 5 presses, even in absence of reward (**Fig. 5f**). This strategy difference is further confirmed by the smaller port-checking probability in LR5 rats (**Fig. 5g)**.

Task-related differences in omission sensitivity were mirrored by distinct DA dynamics in VTA and NAc Core. LI5 rats exhibited a robust decrease in VTA DA neuron activity following reward omission (**Fig. 5j-k**) that was absent in LR5 rats. While VTA DA activity increased in both groups following unexpected reward delivery, this response was greater in LI5 rats (**Fig. 5l-m**). Similarly, NAc Core DA release more strongly decreased after omitted reward in LI5 rats and the response was attenuated in LR5 rats (**Fig. 5n-o**), along with no group differences in response to unexpected reward (**Fig. 5p-q**). Port entry latency after reward omission (**Fig. 5h**) or delivery (**Fig. 5i**) did not differ between groups, ruling out behavioral difference at these events as a source of mesolimbic DA activity discrepancy. These results suggest that LR5 training impairs adaptation to reward omission, by attenuating both negative RPEs following reward omission and positive RPEs following unexpected reward delivery.

### Optogenetic VTA DA neuron stimulation at the time of sequence completion drives automaticity and behavioral chunking

Given that lever retraction paired with reward delivery evokes phasic VTA DA activity and promotes automaticity and behavioral chunking in LR5 rats, we tested whether mimicking this DA activity could replicate the behavior. TH-Cre^+^ rats received unilateral VTA infusions of DIO-ChR2 or DIO-eYFP virus and were implanted with an optic fiber (**Fig. 6a**). We modified the LI5 task (**Fig. 6b**) to incorporate brief optogenetic stimulation (0.5s; 20mW; 20Hz) to simulate the DA burst observed upon sequence completion and lever retraction in the LR5 task. As in the original LI5 task, the lever remained extended until rats retrieved the reward, preventing them from using lever retraction as a reward-predictive cue.

**Figure 6.**
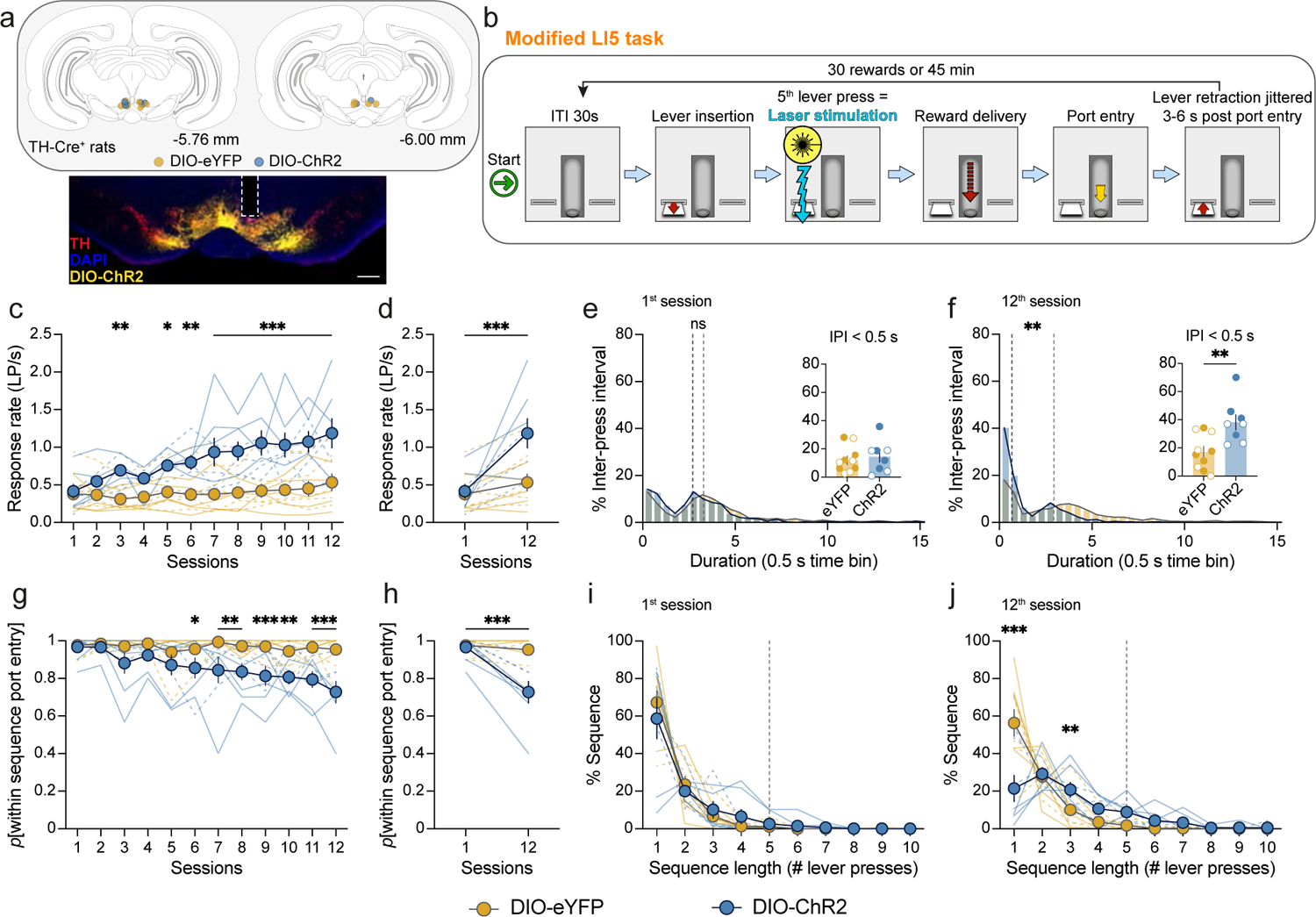
Optogenetic VTA DA neuron stimulation at the time of sequence completion drives automaticity and behavioral chunking. (**a**) Targeting VTA DA neurons with DIO-eYFP or DIO-ChR2 virus in TH-Cre+ rats. Scale bar: 500μm. (**b**) Task design for optogenetic stimulation of VTA DA neurons at sequence completion. (**c**) Response rate in lever press per second (LP/s) within the lever press sequence (two-way RM ANOVA, group effect: F_1,16_ = 15.97, p = 0.001; group x session interaction: F_11,176_ = 4.5, p < 0.001; Student Newman-Keuls post hoc pairwise comparison). (**d**) Response rate on the first versus last session (two-way RM ANOVA, group effect: F_1,16_ = 6.58, p = 0.021; group x session interaction: F_1,16_ = 8.17, p = 0.011; Student Newman-Keuls post hoc pairwise comparison). (**e**) Frequency distribution of inter-press intervals (IPI) for the first session in DIO-eYFP and DIO-ChR2 rats (Mann-Whitney Rank Sum t test median comparison U = 395, p = 0.23). Insert panel: individual proportion of IPI under 0.5 s (t test, no group effect t_16_ = 0.476, p = 0.64). (**f**) Frequency distribution of inter-press intervals (IPI) for the last session in DIO-eYFP and DIO-ChR2 rats (Mann-Whitney Rank Sum t test median comparison U = 274, p = 0.004). Insert panel: individual proportion of IPI under 0.5 s (t test, group effect t_16_ = 3.22, p = 0.005). (**g**) Probability to enter the port within the sequence (within-sequence port entry) (two-way RM ANOVA, group effect: F_1,16_ = 13.84, p = 0.005; group x session interaction: F_11,176_ = 3.95, p < 0.001; Student Newman-Keuls post hoc pairwise comparison). (**h**) Probability of within sequence port entry on the first versus last session in DIO-eYFP and DIO-ChR2 rats (two-way RM ANOVA, group effect: F_1,16_ = 9.41, p = 0.007; group x session interaction: F_1,16_ = 18.06, p < 0.001; Student Newman-Keuls post hoc pairwise comparison). (**i**) Frequency distribution of sequence length during the first training session. The grey dashed line represents the optimal sequence length of 5 lever presses (two-way RM ANOVA, no group effect: F_1,16_ < 1, p = 0.53; no group x sequence length interaction: F_9,144_ < 1, p = 0.77). (**j**) Frequency distribution of sequence length during the last training session. The grey dashed line represents the optimal sequence length of 5 lever presses (two-way RM ANOVA, group effect: F_1,16_ = 5.16, p = 0.037; group x sequence length interaction: F_9,144_ = 8.64, p < 0.001; Student Newman-Keuls post hoc pairwise comparison). Data shown as mean ± SEM, superimposed with individual data point. DIO-eYFP n = 10 (5 males; 5 females); DIO-ChR2 n = 8 (4 males; 4 females). Solid line and filled circles: males; Dashed line and empty circles: females. *p < 0.05; **p < 0.01; ***p < 0.001.

Optogenetic stimulation at sequence completion progressively increased response rates in ChR2 rats across training compared to eYFP rats, also receiving laser presentations (**Fig. 6c-d**). While both groups showed similar IPI distribution in the first session (**Fig. 6e**), by the last session, ChR2 rats exhibited a higher proportion of very short IPIs (**Fig. 6f**). Optogenetic stimulation also reduced port-checking in ChR2 rats (**Fig. 6g**), with all individuals decreasing port entry likelihood between first and last sessions (**Fig. 6h**). Finally, sequence length analysis revealed no group difference during the first session, with mostly single presses before port check (**Fig. 6i**, ∼60%). In the last session, eYFP rats maintained frequent port checking, whereas ChR2 rats made short bouts of 2 or 3 presses before checking the port (**Fig. 6j**), consistent with emerging behavioral chunking.

Although both groups obtained a similar number of rewards, ChR2 rats exceeded the response requirement and retrieved rewards faster than eYFP rats (**Suppl. Fig. 6a-c**). With the parameters used in this experiment (0.5s; 20mW; 20Hz, no cue), ChR2 rats did not show robust intra-cranial self-stimulation (ICSS) in subsequent tests, except for one rat (**Suppl. Fig. 6d-f**). This indicates that VTA DA neuron ICSS does not explain the behavioral effects, as DA-driven responses would be expected to delay, not accelerate, reward retrieval. Importantly, robust ICSS was observed when laser stimulation was increased to 2s (**Suppl. Fig. 6g-i**). In summary, brief unilateral VTA DA neuron activation at sequence completion is sufficient to increase automaticity and, to a lesser extent, behavioral chunking.

## DISCUSSION

We studied how sequence initiation and termination cues differentially influence reward expectation along the sequence and drive habitual and goal-directed strategies. We found that the lever retraction cue marking sequence completion in the LR5 task facilitated habit-like behavior. Conversely, in the LI5 task, in which lever insertion signals trial onset, rats exhibited greater goal-directedness, repeatedly checking for reward between lever presses. These behavioral strategies were associated with distinct mesolimbic DA dynamics, with LR5 rats exhibiting a shift in DA activity from the reward to its predictive cue, while LI5 rats maintained reward-related DA activity with expectation along sequence execution. We also found that behavioral inflexibility in rats trained in the LR5 task was associated with blunted positive and negative RPE-like signals in an omission test. Finally, optogenetic stimulation of VTA DA neurons at sequence termination, in absence of lever retraction cue, was sufficient to increase automaticity and behavioral chunking.

We previously showed that lever cues at sequence boundaries promote habit and behavioral chunking^27,28,33^. These findings have been replicated using other cue types such as auditory tones^30,31,41^. Here, we extended these findings by demonstrating that the lever retraction cue, but not the insertion, is sufficient to promote habit-like behavior. Indeed, LR5-trained rats showed automatic, chunked responses without reward expectation until sequence completion^42,43^. They were also less sensitive to outcome devaluation and negative contingencies in the omission schedule, suggesting reduced behavioral flexibility. These results support the hypothesis that the proximal reward-predictive cue provides response feedback, reducing the need for action-outcome monitoring and promoting habit formation^28,41^. Our prior work^28,44^ showed that rats required to complete five consecutive lever presses without port checking reached LR5-like levels of efficiency, but their behavior relied on internal monitoring and remained goal-directed. Thus, the feedback cue is the critical feature driving the shift toward habitual strategy in the LR5 task.

While DA has been implicated in both sequence^8–10,45^ and habit learning^18,46,47^, these constructs are often studied separately. Yet, in a recent action sequence study, Van Elzelingen et al.^20^ linked shifts in DMS DA release to habit development over extended training, though early learning dynamics were not examined. In contrast, our findings in VTA and NAc show that DA activity is highly dynamic from the very first session, closely tracking changes in reward expectation throughout sequence execution, and reflecting key differences in habitual versus goal-directed strategies. In LR5-trained rats, DA signals rapidly shifted from reward retrieval to the proximal reward predictive stimulus (i.e lever retraction) as previously observed^9,10^. During lever presses, there was no reward expectation, as reflected by the lack of within-sequence port entry and the absence of DA neural activity.

In the LI5 task, DA responses during early training peaked after the first lever press, but with extended training, this peak preceded the press, indicating that rats learned to expect reward after the lever insertion cue and at the lever approach. Additionally, each lever press evoked a phasic VTA DA burst that decayed across the sequence, while no DA release was observed in NAc. This contrasts with prior studies where accumbal DA progressively ramps up during a motivated approach in an action sequence task^10^, decision-making task^48,49^ or when rats have to navigate a maze to obtain a reward^13,50^. In our LI5 task, the absence of ramping DA might be caused by the absence of sensory feedback along the chain of action^51^. Instead, in LI5 rats, inter-press port checking triggered a biphasic VTA DA neuron response, with a brief peak synchronous to the port entry, followed by inhibition of DA activity one second later. This biphasic pattern likely reflects reward expectation when rats entered the port and a negative RPE when reward expectation was violated. The progressive increase in negative RPE magnitude along the sequence suggests that LI5 rats use an internal model of the task, as their reward expectation builds while approaching reward delivery.

We found a training-related decline in DA signals in both VTA and NAc Core in the LR5 task but not in the LI5 task. Similar reductions in cue-evoked DA release have been reported as animals became experts in the task^9,52,53^. This decline is unlikely due to signal bleaching, as DA activity fully recovered following unexpected reward delivery in the omission test. Instead, it may reflect learned expectation of the sequence termination cue^52,53^. Alternatively, with habit formation, termination cue encoding may shift from the mesolimbic to the nigrostriatal pathway^17,19,54–57^. This pathway shift during learning has been suggested as a neural basis for the development of automaticity and the transition from goal-directed to habitual control^18,19,54^.

Dickinson et al.^58^ first showed that extensive reinforcement history and habit learning drives behavioral persistence under an omission schedule. Accordingly, training in the LI5 and LR5 tasks lead to distinct behavioral and dopaminergic adaptations during the omission test. LI5-trained rats maintained goal-directed control and were able to flexibly reduce their response in absence of reward. Their neural signature showed negative RPE following reward omission and positive RPE following unexpected reward delivery. Conversely, LR5 rats’ resistance to reward omission was associated with attenuated positive RPE and blunted negative RPE, suggesting reduced sensitivity to negative contingency. Studies have shown that DA responses to conditioned reward cues and reward omission scale with uncertainty of reinforcement^6,59^. Under high uncertainty, VTA DA neurons are activated by reward omission, potentially driving adaptive learning and behavioral switching to obtain reward^60^. After training under reward uncertainty in the LI5 task, repeated negative RPEs after reward omission likely drives behavioral extinction and adaptation to the reversal in instrumental contingencies. In contrast, when reward delivery is predicted with certainty by the lever retraction in the LR5 task, rats may persist to respond, leading to habitual behavior that is less sensitive to outcome omission and relies more on cues. These results suggest that the lack of negative RPE signal is linked to the reduced behavioral flexibility that characterizes habits.

Dopamine-dependent synaptic plasticity in corticostriatal circuits is essential for habit learning^18,47^. In the striatum, sensorimotor events such as lever presses, trigger synaptic activity that leaves behind eligibility traces lasting approximatively 1-2 seconds^36,61^. If a reward is delivered during this window, DA signaling can induce synaptic plasticity, facilitating credit assignment between an action and its outcome^36,61–63^. Our lab showed that VTA DA neuron ICSS in an action sequence task results in the backpropagation of the reinforcement signal, promoting sequence completion^8^. Similarly, another study showed that optogenetic stimulation of the DMS at the first lever press promotes habit^20^. Interestingly, both studies involved seeking-taking chain schedules in which pressing on a lever triggered extension of another lever and pressing on this second lever resulted in reward delivery. This type of task may facilitate the backpropagation of eligibility traces from proximal cues to distal cues. Here, the sequence termination cue elicited a transient DA response as rats learned to complete the LR5 sequence. Mimicking this DA signal with optogenetics, without the actual lever retraction cue, promoted partial automaticity and behavioral chunking. A possible mechanism is that optogenetic DA stimulation time-locked to the final lever press facilitates credit assignment between sequence completion and the reward. With repetition, this may support the backward spread of credit to earlier action in the sequence, ultimately reinforcing the sequence as a whole unit^61^.

Notably, DA-stimulated rats often exceeded the required number of lever presses to receive the reward. While this might resemble ICSS, the brief (0.5s) VTA stimulation used here lacks the strong reinforcing effect seen with longer (1s) stimulation durations reported in other studies^2,3,64,65^ (but see^66^). Despite this, ChR2 rats remained engaged in the task, as evidenced by the number of rewards obtained and a short retrieval latency. Thus, the additional lever presses likely indicate the difficulty in inhibiting a prepotent behavior due to increased speed in sequence execution. Likewise, LR5 trained rats often pressed on the lever while retracting, leading to sequences of 6 or 7 lever presses (**Fig. 1**).

To conclude, we found sequence initiation and termination cues differentially affect reward expectation along the sequence and drive distinct response strategies that are encoded by specific mesolimbic DA patterns. The goal-directed strategy was associated with recurrent and increasing DA-encoded reward expectation throughout the sequence whereas the habitual strategy was marked by DA signals confined to the sequence termination and attenuated RPEs following expectation violations. These findings underscore the crucial role of mesolimbic DA signals at sequence termination in assigning credit to the whole action sequence to drive automated and habitual behavior.

## Supporting information

Table 1_Satistics sex differences

## ACKNOWLEDGEMENTS

The authors thank all members of the Janak lab for their helpful discussions on the experimental findings.

## FUNDING AND DISCLOSURES

This work was supported by NIH grant R01DA035943 (PHJ) and IRESP grant AAPSPA2021-V1-07 (NT). The authors declare no conflict of interest.

## AUTHOR CONTRIBUTIONS

RM, YV and PHJ designed the experiments; RM, YV, collected the data with technical assistance from JZ, YC and HP; RM, YV analyzed the data with assistance from YC; PHJ and NT provided financial support; RM, YV and PHJ interpreted the data and wrote the manuscript. RM, YC, JZ, HP, NT, PHJ and YV provided inputs, made edits and approved the final version of the manuscript.

## Data and code availability

Data and python code used for analysis and figures have been deposited at G-node and is publicly available as of the date of publication. Any additional information required to reanalyze the data reported in this paper is available from the lead contact upon request.

## Notes

### Competing Interest Statement

The authors have declared no competing interest.

### Summary of Updates

New supplemental figure (Supply. Fig. 3) on dopamine activity during within sequence port entry. Overall manuscript length shorten Discussion reshaped

