## Supplemental materials for "Sequence termination cues drive habits via dopamine-mediated credit assignment"

#### CORRESPONDENCE

#### SUPPLEMENTAL FIGURES

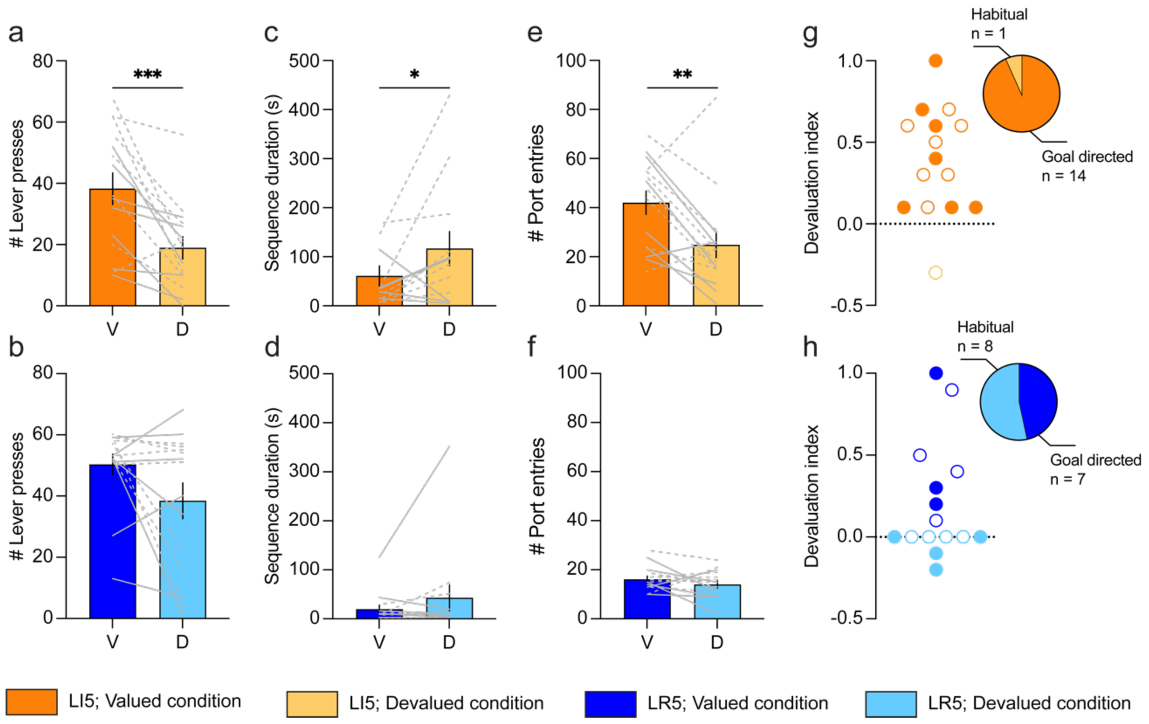


**Supplemental Figure 1: LR5 task training favors insensitivity to satiety-induced outcome devaluation in a subset of individuals**. (**a**) Number of lever presses performed by LI5 rats in the valued (V) and devalued (D) conditions (one-way RM ANOVA, devaluation effect: $F_{1,14}$ = 18.7, p < 0.001; Student Newman-Keuls post-hoc pairwise comparison). (**b**) Same as (**a**) in the LR5 task (one-way RM ANOVA, devaluation effect: $F_{1,14}$ = 4.32, p = 0.056). (**c**) Time to complete the lever press sequence for LI5 rats in the valued (V) and devalued (D) conditions (one-way RM ANOVA, devaluation effect: $F_{1,12}$ = 5.23, p = 0.041). (**d**) same as (**c**) in the LR5 task (Friedman RM ANOVA, no devaluation effect: $\chi^2$ = 0.692, p = 0.581). (**e**) Number of port entries performed by LI5 rats in the valued (V) and devalued (D) conditions (one-way RM ANOVA, devaluation effect: $F_{1,14}$ = 12.2, p = 0.004; Student Newman-Keuls post hoc pairwise comparison). (**f**) same as (**e**) in the LR5 task (one-way RM ANOVA, no devaluation effect: $F_{1,14}$ = 1.53, p = 0.236). (**g - h**) Different proportions of goal-directed or habitual rats trained in the LI5 task (**g**) or LR5 task (**h**) ($\chi^2$ = 6.2, p = 0.013). LI5 n = 15 (7 males; 8 females); LR5 n = 15 (7 males; 8 females). Data are shown as means ± SEM, superimposed with individual data point. Males: solid lines and filled circles; Females: dashed lines and empty circles. **p < 0.01; ***p < 0.001.

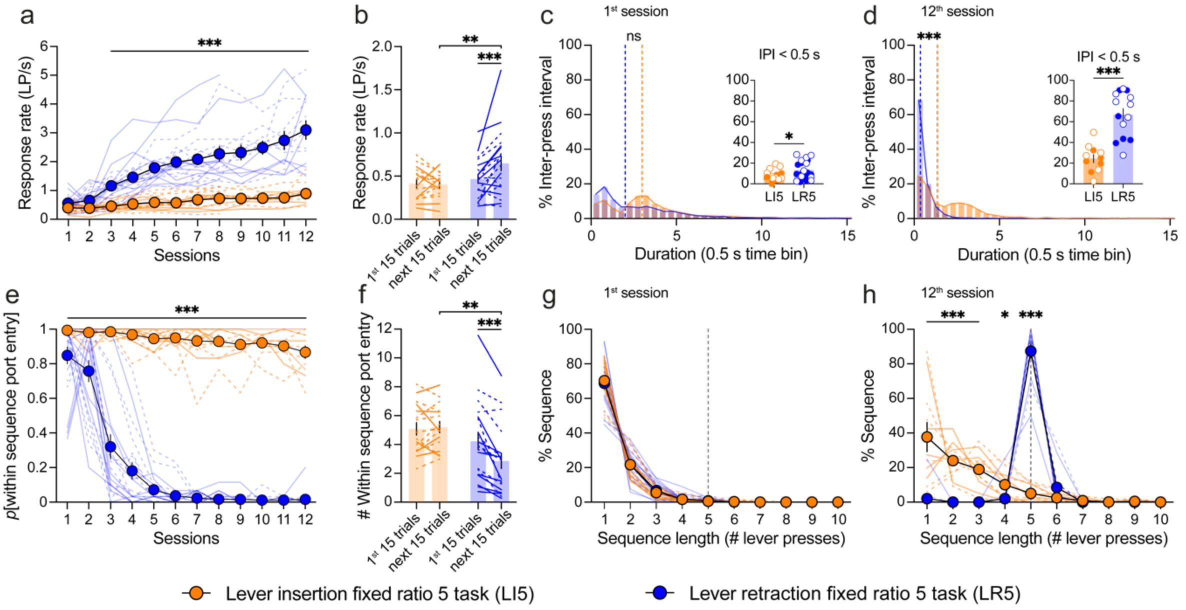


**Supplemental Figure 2: The lever retraction cue, but not the lever insertion cue, favors action automaticity and behavioral chunking during photometry recording (VTA and NAc).** (**a**) Response rate in lever press per second (LP/s), within the lever press sequence (three-way RM ANOVA, task effect: $F_{1,21} = 50.92$, $p < 0.001$; task x session interaction: $F_{11,231} = 10.32$, $p < 0.001$; no task x session x region interaction: $F_{11,231} < 1$, $p = 0.551$; Student Newman-Keuls post-hoc pairwise comparison). (**b**) Response rate for the first 15 trials versus next 15 trials of LI5/LR5 training (three-way RM ANOVA, task effect: $F_{1,32} = 4.75$, $p = 0.037$; task x trials interaction: $F_{1,32} = 7.13$, $p = 0.012$; no task x session x region interaction: $F_{1,32} = 1.05$, $p = 0.314$; Student Newman-Keuls post-hoc pairwise comparison). (**c**) Frequency distribution of inter-press intervals (IPI) for the 1$^{st}$ LI5/LR5 session (Mann-Whitney Rank Sum t-test median comparison U = 473, $p = 0.921$). Insert panel: individual proportion of IPI under 0.5 s (two-way ANOVA, task effect: $F_{1,32} = 4.39$, $p = 0.044$; no task x region interaction: $F_{1,32} = 1.06$, $p = 0.310$; Student Newman-Keuls post-hoc pairwise comparison). (**d**) Frequency distribution of IPI during the 12$^{th}$ LI5/LR5 session (Mann-Whitney Rank Sum t-test median comparison U = 185, $p < 0.001$). Insert panel: individual proportion of IPI under 0.5 s (two-way ANOVA, task effect: $F_{1,21} = 31.04$, $p < 0.001$; no task x region interaction: $F_{1,21} = 1.37$, $p = 0.255$). (**e**) Probability to enter the port within the lever press sequence (within-sequence port entry) (three-way RM ANOVA, task effect: $F_{1,21} = 1136.72$, $p < 0.001$; task x session interaction: $F_{11,231} = 38.76$, $p < 0.001$; task x session x region interaction: $F_{11,231} = 3.252$, $p < 0.001$; Student Newman-Keuls post-hoc pairwise comparison). (**f**) Number of within-sequence port-entry for the first 15 trials versus next 15 trials of the LI5/LR5 tasks (three-way RM ANOVA, task effect: $F_{1,32} = 6.33$, $p = 0.017$; task x trials interaction: $F_{1,32} = 19.60$, $p < 0.001$; no task x session x region interaction: $F_{1,32} < 1$, $p = 0.738$; Student Newman-Keuls post hoc pairwise comparison). (**g**) Frequency distribution of sequence length during the first training session in the LI5/LR5 tasks. The grey dashed line represents the optimal sequence length of 5 lever presses (three-way RM ANOVA, no task x sequence length interaction: $F_{9,288} < 1$, $p = 0.98$; no task x sequence length x region interaction $F_{9,288} < 1$, $p = 0.99$). (**h**) Frequency distribution of sequence length during the last training session in the LI5/LR5 tasks. The grey dashed line represents the optimal sequence length of 5 lever presses (three-way RM ANOVA, task effect: $F_{9,189} = 84$, $p < 0.001$; task x sequence length interaction: $F_{9,189} = 73.90$, $p < 0.001$; task x sequence length x region interaction $F_{9,189} = 2.422$, $p = 0.013$; Student Newman-Keuls post hoc pairwise comparison). Data are shown as mean ± SEM, superimposed with individual data point. LI5: n = 11-16 (5-6 males; 6-11 females); LR5: n = 14-20 (6-10 males; 8-10 females). Regions: VTA: n = 18 (4-10 LR5; 3-8 LI5); NAc: n = 18 (10 LR5; 8 LI5). Solid lines and filled circles: males; dashed lines and empty circles: females. *$p < 0.05$; **$p < 0.01$; ***$p < 0.001$.

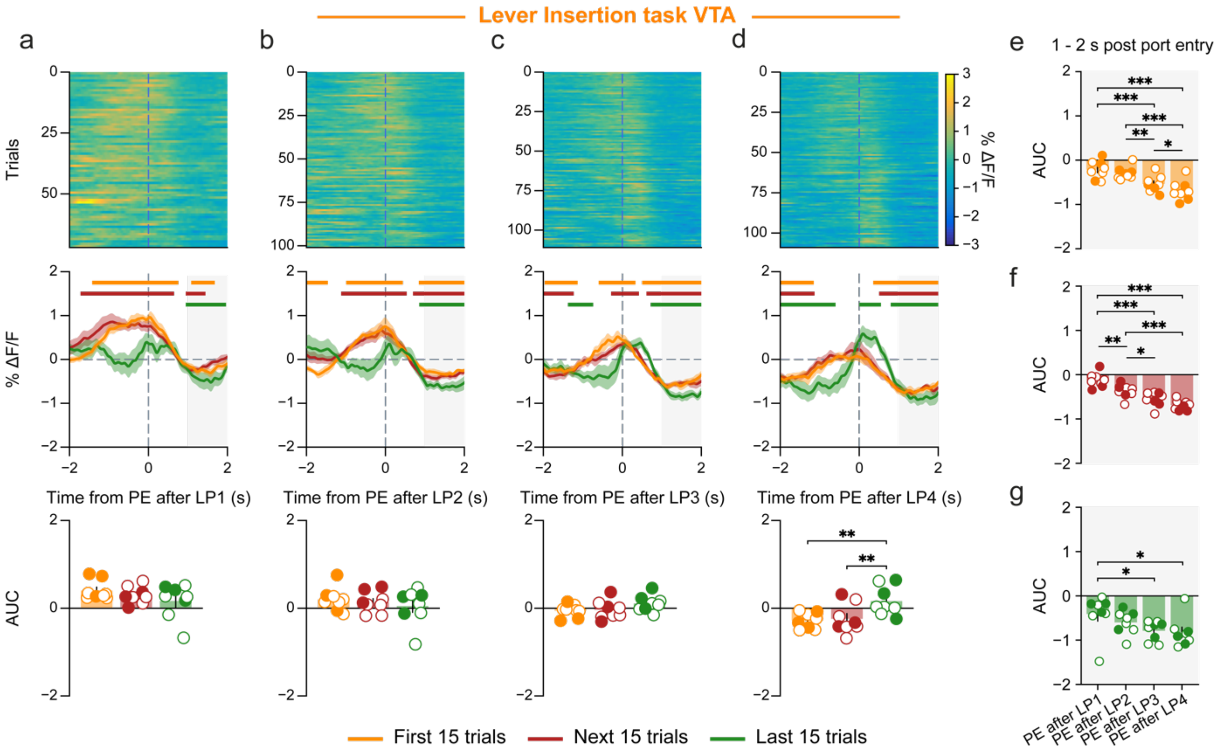


**Supplemental Figure 3: Patterns of VTA DA neuron activity during port entry following each lever press of the sequences in the LI5 task**. (**a-d**) First port entry following the first four lever presses of the sequence. The upper panel heatmaps illustrate trial-by-trial patterns of dopamine neuron activity, averaged across rat. The middle panel represents phasic dopamine release averaged across rats and trials, delineated for the initial 15 trials (orange line), the subsequent 15 trials (red line), and the final 15 trials (green line) of the training period. The lower panel displays the area under the curves (AUCs) for the 0 to 1 second time range following the event. (**a**) Port entry following the 1$^{st}$ lever press (one-way RM ANOVA, no trial effect: $F_{2,14} = 1.22$, $p = 0.324$). (**b**) Port entry following the 2$^{nd}$ lever press (one-way RM ANOVA, no trial effect: $F_{2,14} < 1$, $p = 0.496$). (**c**) Port entry following the 3$^{rd}$ lever press (one-way RM ANOVA, no trial effect: $F_{2,14} = 1.60$, $p = 0.236$). (**d**) Port entry following the 4$^{th}$ lever press (one-way RM ANOVA, trial effect: $F_{2,14} = 7.43$, $p = 0.006$; Student Newman-Keuls post-hoc pairwise comparison). (**e-g**) Area under the curves (AUCs) for the 1 to 2 second window following the port entry after the first four lever presses of the sequence in the LI5 task (grey shaded area on PSTH), measured across the three trial periods (two-way RM ANOVA, trial effect: $F_{2,14} = 7.13$, $p = 0.007$; port entry effect: $F_{3,21} = 20.89$, $p < 0.001$; no port entry x trial interaction $F_{6,42} < 1$, $p = 0.782$). (**e**) First 15 trials (one-way RM ANOVA, port entry effect: $F_{3,21} = 20.353$, $p < 0.001$; Student Newman-Keuls post-hoc pairwise comparison). (**f**) Next 15 trials (one-way RM ANOVA, port entry effect: $F_{3,21} = 23.09$, $p < 0.001$; Student Newman-Keuls post-hoc pairwise comparison). (**g**) Final 15 trials (one-way RM ANOVA, port entry effect: $F_{3,21} = 3.564$, $p = 0.032$; Student Newman-Keuls post-hoc pairwise comparison). Data are shown as means ± SEM, superimposed with individual data point. LI5 n = 8 (3 males; 5 females). Horizontal bars indicate significant deviation from baseline at 99% confidence level. Filled circles: males; Empty circles: females. *$p < 0.05$; **$p < 0.01$; ***$p < 0.001$.

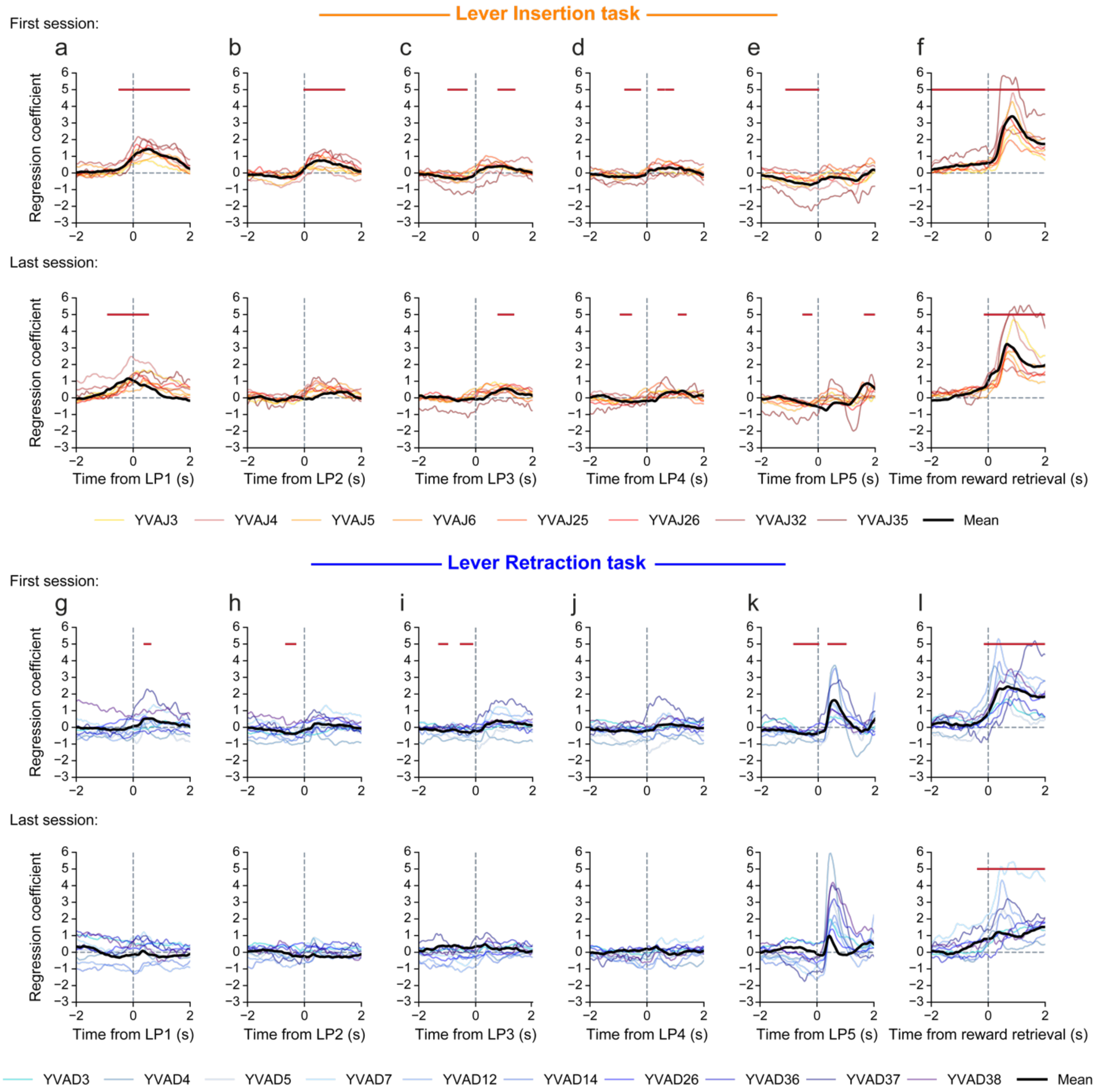


**Supplemental Figure 4: Deconvolution analysis. (a-b)** Mean regression coefficients around each lever press event and reward retrieval during the first (**a**) and last (**b**) training session in the lever insertion task. (**c-d**) Mean regression coefficients around each lever press event and reward retrieval during the first (**c**) and last (**d**) training session in the lever retraction task. Colored lines represent individual rats. Black lines represent the average regression coefficient across rats. Horizontal red bars indicate significant deviation from baseline at 99% confidence level.

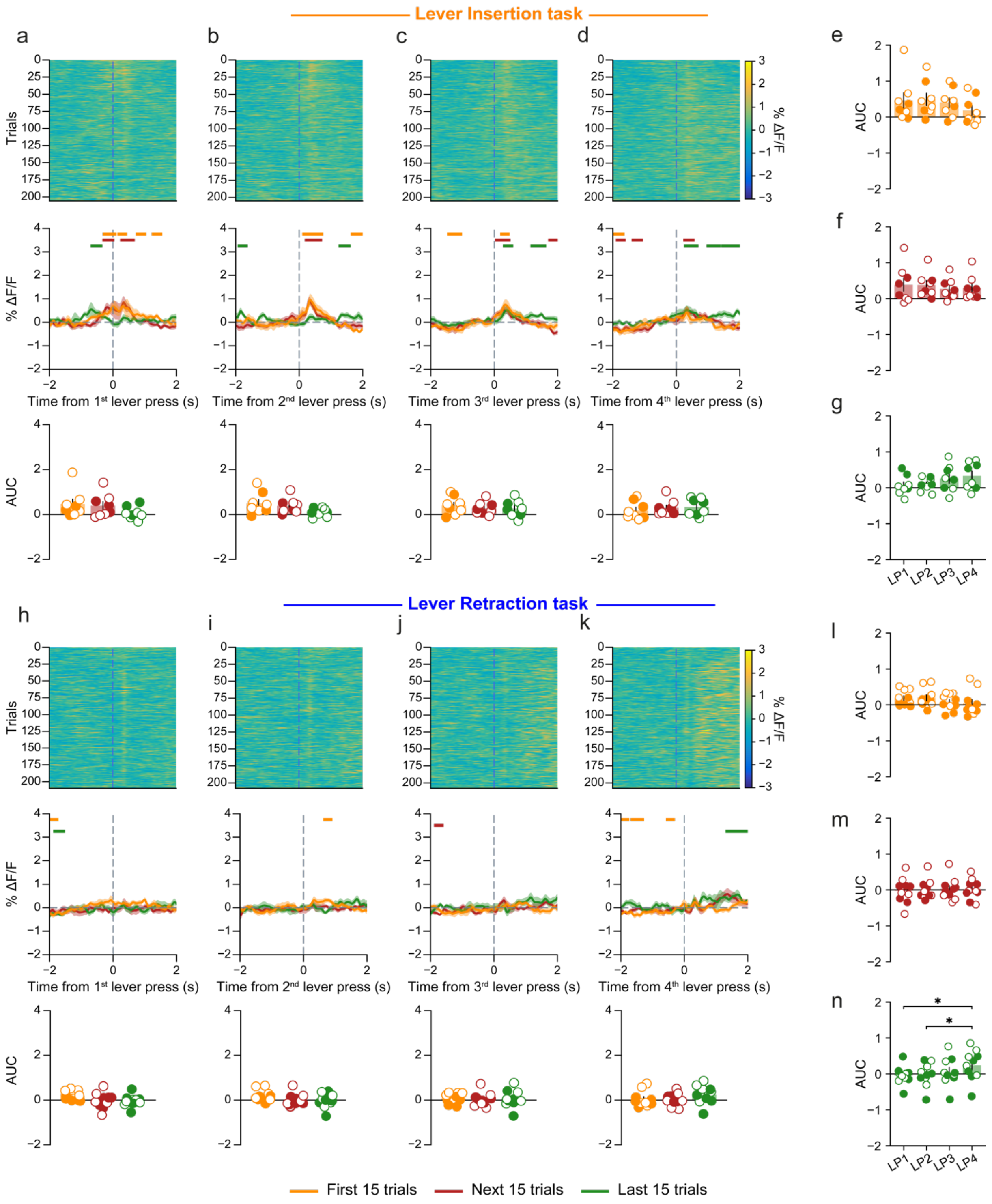


**Supplemental Figure 5: Distinct patterns of dopamine release in the NAc Core during lever press sequences in the two tasks**. (**a-d**) Sequence of the first four lever presses for the LI5 task. The upper panel heatmaps illustrate trial-by-trial patterns of dopamine release, averaged across rat. The middle panel represents phasic dopamine release averaged across rats and trials, delineated for the initial 15 trials (orange line), the subsequent 15 trials (red line), and the final 15 trials (green line) of the training period. The lower panel displays the area under the curves (AUCs) for the 0 to 1 second time range following the event. (**a**) 1$^{st}$ lever press (LP1; one-way RM ANOVA, no trial effect: $F_{2,14}$ = 2.15, p = 0.154). (**b**) 2$^{nd}$ lever press (LP2; one-way RM ANOVA, no trial effect: $F_{2,14}$ = 3.36, p = 0.064). (**c**) 3$^{rd}$ lever press (LP3; one-way RM ANOVA, no trial effect: $F_{2,14}$ < 1, p = 0.635). (**d**) 4$^{th}$ lever press (LP4; one-way RM

ANOVA, no trial effect: $F_{2,14} < 1$, $p = 0.65$). (**e-g**) Area under the curves (AUCs) for the 0 to 1 s window following the first four lever presses of the sequence in the LI5 task, measured across the three trial periods (two-way RM ANOVA, no trial effect: $F_{2,14} = 0.884$, $p = 0.435$; no lever press effect: $F_{3,21} = 0.204$, $p = 0.892$; lever press x trial interaction $F_{6,42} = 4.296$, $p = 0.002$). (**e**) First 15 trials (one-way RM ANOVA, no lever press effect: $F_{3,21} = 2.731$, $p = 0.07$). (**f**) Next 15 trials (one-way RM ANOVA, no lever press effect: $F_{3,21} = 1.45$, $p = 0.258$). (**g**) Final 15 trials (one-way RM ANOVA, no lever press effect: $F_{3,21} = 2.56$, $p = 0.08$). (**h-k**) same as (**a-d**) for the LR5 task. (**h**) 1$^{st}$ lever press (LP1; one-way RM ANOVA, no trial effect: $F_{2,18} = 2.35$, $p = 0.12$). (**i**) 2$^{nd}$ lever press (LP2; one-way RM ANOVA, no trial effect: $F_{2,18} = 2.47$, $p = 0.11$). (**j**) 3$^{rd}$ lever press (LP3; one-way RM ANOVA, no trial effect: $F_{2,18} < 1$, $p = 0.91$). (**k**) 4$^{th}$ lever press (LP4; one-way RM ANOVA, no trial effect: $F_{2,18} = 2.32$, $p = 0.13$). (**l-n**) same as (**e-g**) for the LR5 task (two-way RM ANOVA, no trial effect: $F_{2,18} = 0.785$, $p = 0.471$; no lever press effect: $F_{3,27} = 618$, $p = 0.609$; lever press x trial interaction $F_{6,54} = 3.62$, $p = 0.04$). (**l**) First 15 trials (one-way RM ANOVA, no lever press effect: $F_{3,27} = 2.12$, $p = 0.12$). (**m**) Next 15 trials (one-way RM ANOVA, no lever press effect: $F_{3,27} < 1$, $p = 0.84$). (**n**) Final 15 trials (one-way RM ANOVA, lever press effect: $F_{3,27} = 3.57$, $p = 0.03$; Student Newman-Keuls post-hoc pairwise comparison). Data are shown as means ± SEM, superimposed with individual data point. LI5 n = 8 (3 males; 5 females); LR5 n = 10 (5 males; 5 females). Horizontal bars indicate significant deviation from baseline at 99% confidence level. Filled circles: males; Empty circles: females. *$p < 0.05$.

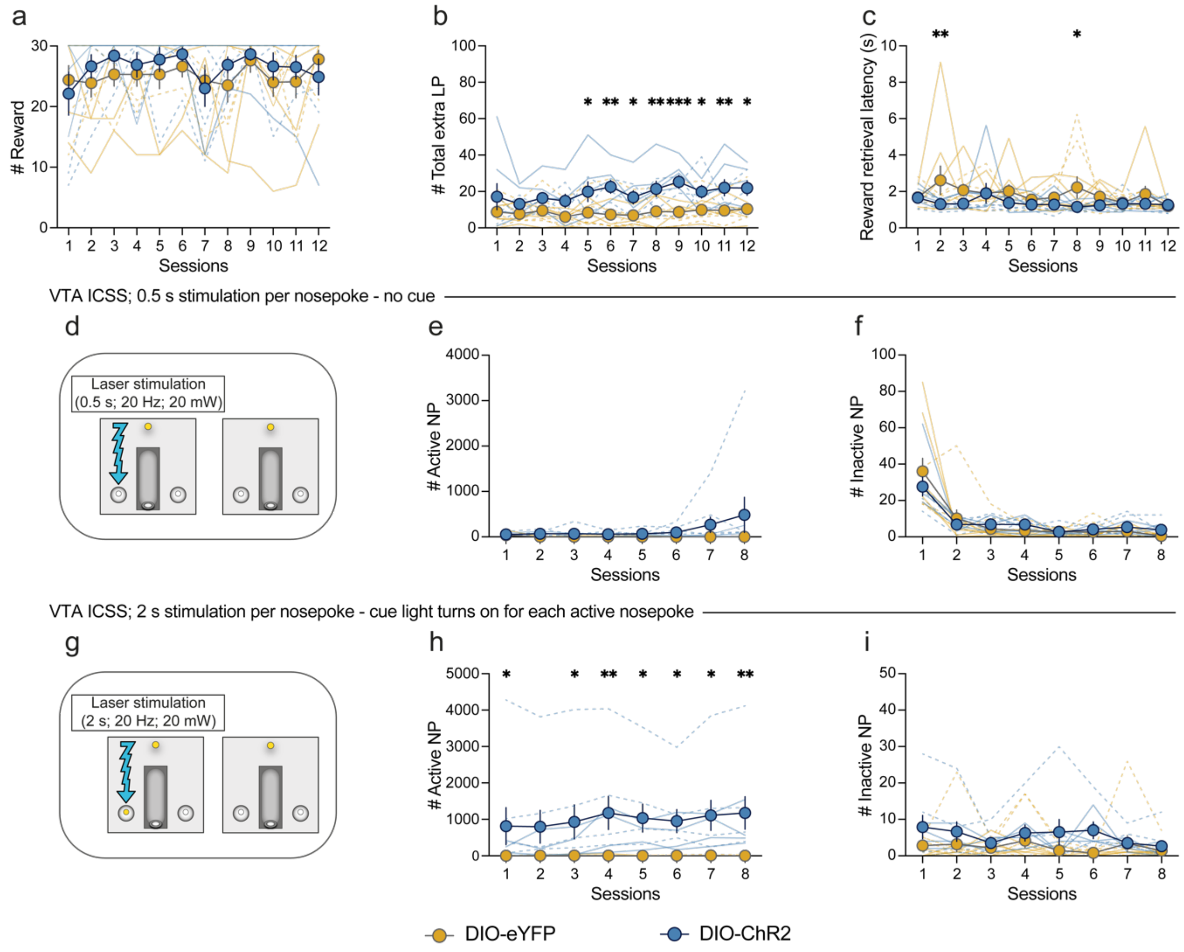


**Supplemental Figure 6: VTA DA optogenetic stimulation at the time of sequence completion has moderate reinforcing effect.** (**a**) Number of rewards obtained per session (two-way RM ANOVA, no group effect: $F_{1,16} < 1$, p = 0.579; no group x session interaction: $F_{11,176} < 1$, p = 0.626). (**b**) Number of extra lever press emitted after sequence completion (two-way RM ANOVA, group effect: $F_{1,16} = 9.187$, p = 0.008; no group x session interaction: $F_{11,176} = 1.24$, p = 0.263; Student Newman-Keuls post hoc pairwise comparison). (**c**) Reward retrieval latency (two-way RM ANOVA, group effect: $F_{1,16} = 5.67$, p = 0.03; no group x session interaction: $F_{11,176} = 1.02$, p = 0.434; Student Newman-Keuls post hoc pairwise comparison). (**d**) Schematic representation of VTA ICSS with the stimulation parameters used in the main experiment (0.5 s; 20 mW; 20 Hz, in absence of cue). (**e**) Number of active nose pokes per session (two-way RM ANOVA, no group effect: $F_{1,16} = 3.776$, p = 0.070; no group x session interaction: $F_{7,112} = 1.645$, p = 0.13). (**f**) Number of inactive nose pokes per session (two-way RM ANOVA, no group effect: $F_{1,16} < 1$, p = 0.926; no session x group interaction: $F_{7,112} = 1.05$, p = 0.402). (**g**) Schematic representation of VTA ICSS with 2 s stimulation and cue light during active nose pokes. (**h**) Number of active nose pokes per session (two-way RM ANOVA, group effect: $F_{1,16} = 6.91$, p = 0.018; group x session interaction: $F_{7,112} = 3.64$, p = 0.001; Student Newman-Keuls post hoc pairwise comparison). (**i**) Number of inactive nose pokes per session (two-way RM ANOVA, no group effect: $F_{1,16} = 3.01$, p = 0.10; no group x session interaction: $F_{7,112} = 1.44$, p = 0.196). Data are shown as means ± SEM, superimposed with individual data point. DIO-eYFP n = 10 (5 males; 5 females); DIO-ChR2 n = 8 (4 males; 4 females). Solid line: males; Dash line: females. *p < 0.05; **p < 0.01; ***p < 0.001.

### SUPPLEMENTAL MATERIALS & METHODS

#### Subjects

Wild-type (WT) (n = 48; 23 males and 25 females) or TH-Cre$^+$ Long-Evans rats (n = 36; 17 males and 19 females) were used in this study. These rats express Cre recombinase under the control of the tyrosine hydroxylase (TH) promoter[64]. Rats were individually housed under a 12 h/light/dark cycle (lights ON at 7 am). They were food restricted to 90% of free feeding weight, starting one week before training, but had *ad libitum* access to water throughout the experiment. All rats were 8 to 10 weeks old at the beginning of the experiment. These studies were carried out in accordance with the recommendations of the Guide for the Care and Use of Laboratory Animals (Institute of Instrumental Training Laboratory Animal Resources, Commission of Life Sciences, National Research Council, 1996). The protocol was approved by the animal care and use committee of Johns Hopkins University.

#### Behavioral training

##### *Apparatus*

The apparatus consisted of operant chambers (Med associates) each equipped with a retractable lever on the left side of the right wall. The reward (0.1mL of 20% sucrose) was delivered over 3 s in a magazine equipped with an infra-red beam, and located next to the lever. The house-light, located on the back wall of the operant chamber remained illuminated for the length of the session.

##### Initial training

Rats experienced one magazine training session in which 30 sucrose rewards were delivered under a random interval 60 s schedule. The following days, they were trained to press the lever to earn reward on a fixed-ratio 1 (FR1) schedule. Sessions were limited to 1 hour or 30 reward deliveries, whichever occurred first. Rats were trained one session per day, for 3 to 5 sessions, until they earned the maximum 30 rewards in less than an hour. Rats were then trained on a FR5 schedule (five responses per reward) for 5 additional sessions, limited to 30 min or 30 reward deliveries, whichever occurred first. During initial instrumental training, the lever was continuously presented and no cue indicated when reward was delivered.

##### *LI5 and LR5 task design*

The lever insertion fixed-ratio 5 (LI5) task and the lever retraction fixed-ratio 5 (LR5) task were designed such that only one of the lever cues (lever insertion or retraction, respectively) is relevant to signal reward while the other cue, is made irrelevant. Both groups of rats were trained in parallel in the LI5 task or LR5 task. Two rats were excluded from analyses because they omitted the reward more than 70% of the trials on average across the 12 sessions, leaving a total of 30 rats (LI5: n=15, 8 females, 7 males; LR5: n=15, 8 females, 7 males).

###### *LI5 task*

As detailed in **Fig 1a**, each trial begun with a 30 s inter-trial interval prior to the insertion of the lever in the cage. Rats were then required to press the lever five times to receive the reward. The fifth lever press triggered reward delivery, which was not signaled by the lever retraction cue. To ensure that rats did not pay attention to the retraction of the lever, this event occurred during reward consumption, at a random interval ranging from 3 to 6 s after rats' entry into the magazine or at the exit of the magazine, whichever occurred first. The session ended after 45 min or 30 reward deliveries. For each trial, the lever remained extended until the rats completed the ratio and entered the magazine to consume the sucrose.

###### *LR5 task*

As detailed in **Fig 1b**, each trial begun with the insertion of the lever in the cage. The completion of the FR5 requirement led to the retraction of the lever which signaled reward delivery, occurring at the fifth lever press. To ensure that rats did not pay attention to the insertion of the lever, this event occurred at a random interval ranging from 3 to 6 s after the rats entered the magazine to consume the sucrose or at magazine exit, whichever occurred first. The session ended after 30 min or 30 reward deliveries. For each trial, the lever remained extended until completion of the ratio.

##### *Outcome devaluation by sensory-specific satiety*

In the first experiment, habit learning was assessed after 12 training sessions using the outcome devaluation test. Each rat received 2 days of testing, separated by one reinforced training session. Rats were given 1 hour free access to their training reward (sucrose 20%; devalued condition) or to a control reward, which never served as a reinforcer (grain-based pellet; valued condition). Pre-feeding occurred in feeding cages in the experimental room. Immediately after pre-feeding, rats were placed in the operant chambers for a test session conducted under extinction with the same LI5/LR5 task design. Test sessions were limited to 10 trials or 10 minutes of lever presentation, whichever occurred first. Experimental valued/devalued conditions were reversed for the second test session and the order of conditions was counterbalanced across rats.

##### *Omission test*

In the second and third experiments involving fiber photometry recording in the VTA and NAc, habit learning was assessed after 8 or 12 training sessions by testing response sensitivity to an omission schedule. Fiber photometry recording was conducted during this test session. As illustrated in **Fig 5a-b** the omission schedule consisted on a single test session in which animals had to refrain from pressing the lever for 20 s in order to obtain the reward. In contrast, completion of the FR5 resulted in omission of the reward. Every lever press reset the 20 s timer. The fundamental structures of the LI5 and LR5 tasks were maintained: for rats trained in the LI5 task, lever insertion indicated the initiation of a trial, while for those trained in the LR5 task, lever retraction denoted the fulfillment of the response requirement. The session only ended after 30 rewards were delivered, with no time limit.

#### Surgical procedure

Surgeries for viral infusions and optic fiber implants were carried out as previously described[3,65]. Rats were anesthetized with 5% isoflurane and placed in a stereotaxic frame, after which anesthesia was maintained at 1–3%. Rats were administered carprofen anesthetic (5 mg/kg), and cefazolin antibiotic (70 mg/kg) subcutaneously. The top of the skull was exposed and holes were drilled for viral infusion, optic fiber implant, and four skull screws. Viral injections were made using a microsyringe pump at a rate of 0.1 µl/min. Syringes were left in place for 5 min, then raised 200 µm dorsal to the injection site, left in place for another 10 min, then removed slowly. Implants were secured to the skull with dental cement. Optogenetic manipulations commenced at least 4 weeks after surgery. A total of 5 to 6 weeks was allowed for virus expression prior to photometry recording.

##### *Fiber photometry*

For fiber photometry recording of VTA dopamine neurons bulk calcium imaging, 1 µl Cre-dependent GCaMP6f (AAVDJ-EF1a-DIO-GCaMP6f; titer $5.25 \times 10^{12}$ particles/ml, Stanford University) was infused unilaterally into the VTA (AP -5.8 mm, ML ±0.7 mm, DV -7.9 mm relative to Bregma) of TH-Cre$^+$ rats. In order to avoid residual florescence recording from SNc, low-auto-fluorescence optic fibers (400 µm diameter; 0.57 numerical aperture, Doric Lenses) were inserted with a ±10° angle, directed toward the VTA midline, just dorsal to the injection site (AP -5.8 mm, ML ±1.7 mm, DV -7.4 mm relative to Bregma). Fiber photometry recording of dopamine signaling in the NAc was achieved via infusion of the dopamine sensor dLight1.2 virus (pAAV5-hSyn-dLight1.2: titer 5-5.75 $\times 10^{12}$ particles/ml, addgene) in the NAc Core (AP +1.8 mm, ML ±1.7 mm, DV -7 mm relative to Bregma) of WT rats. Optic fibers in the NAc Core were implanted at AP +1.8 mm, ML ±1.7 mm, DV -6.8 mm relative to Bregma.

##### *Optogenetics*

AAV5-Ef1a-DIO-ChR2-eYFP (titer 4–4.2 × $10^{12}$ particles/ml, University of North Carolina) or AAV5-EF1a-DIO-eYFP (titer 4–6 × $10^{12}$ particles/ml, University of North Carolina) were infused unilaterally (0.7 µl at each target site, for a total of 2.8 µl per rat, as previously described [2,3]) at the following coordinates from Bregma for targeting VTA cell bodies: AP -6.2 and -5.4 mm, ML ±0.7 mm, DV -8.5 and -7.5. Custom-made optic fiber implants (300-µm glass diameter) were inserted unilaterally in the VTA, just above and between viral injection sites at the following coordinates relative to Bregma: AP -5.8 mm, ML ±0.7 mm, DV -7.5.

#### Fiber photometry recording

We assessed VTA dopamine neuron activity in TH-Cre$^+$ rats during training in the LI5 task (n = 8; 5 females, 3 males) or the LR5 task (n = 10; 5 females, 5 males). Dopamine release in the NAc Core was assessed in WT rats trained in the LI5 task (n = 8, 5 females, 3 males) or the LR5 task (n = 10, n= 5 females, 5 males). For the group of rat recorded in the VTA, 3 LI5 trained and 6 LR5-trained did not experienced a jittered insertion of the lever in the cage. Instead, for the LI5 trained rats, the lever was retracted in the cage 0.5 s after the port entry following ratio completion. For the LR5-trained rats, the lever was inserted in the cage 1.5 s after the port entry following ratio completion.

Operant chambers were equipped with a real-time fiber photometry recording system (Tucker-Davis Technologies). A fluorescence mini-cube (Doric Lenses) was utilized to transmit light streams from a 465 nm LED modulated at 211 Hz and a 405 nm LED modulated at 531 Hz. The 465 nm LED passed through a GFP excitation filter, while the 405 nm LED passed through a 405 nm bandpass filter. The power of the LEDs was set at approximately 100 µW. These light sources were connected to an optic fiber implanted in the rat's brain. Power output (~30 µW and ~70 µW for the 405 nm and the 465 nm channel respectively) was daily measured at the tip of the patch cord, before and after each rat's session to ensure a stable power across rats and recording days.
Both GCaMP6f and dLight fluorescence emitted by neurons below the fiber tip were captured by the mini-cube and passed through a GFP emission filter. The fluorescence signal was then amplified and focused onto a high sensitivity photoreceiver (Newport, Model 2151). To account for bleaching and movement artifacts, the brightness produced by the 465 nm excitation, which stimulates calcium-dependent GCaMP6f fluorescence, was demodulated and compared to the isosbestic 405 nm excitation, which stimulates GCaMP6f or dLight in a calcium and dopamine-independent manner.
Rats received initial operant training under the FR1 schedule before surgery Five to six weeks after virus infusion, rats were given two FR1 reminder sessions to acclimate rats to patch cord tethering and ensure they were able to move easily. Photometry recording was conducted throughout 8 training sessions in the LI5 and LR5 tasks and during the omission test.

#### Optogenetic stimulation

##### *Optogenetic stimulation in the modified-LI5 task*

Illumination was provided by 473 nm lasers (OptoEngine), adjusted to read 20 mW from the tip of the patch cord at constant illumination. Light power was measured before and after every behavioral session to ensure that laser power was constant within session. Black tape was wrapped around the ceramic sleeve used to connect the patch cord and the implanted optic fiber to block visible light transmission that could be used as a cue during the task. A total of 8 ChR2 rats (4 females, 4 males) and 10 eYFP (5 females, 5 males) were run in this experiment. As shown in **Fig 6b**, the LI5 task was modified to include a brief laser stimulation (0.5 s; 20 mW; 20 Hz, 5 ms pulse duration) at the 5$^{th}$ lever press. Rats were trained for 12 consecutive sessions that terminated after 45 minutes or 30 rewards obtained, whichever occurred first.

##### *Intra-cranial self-stimulation (ICSS)*

At the end of the optogenetic experiment, all rats were tested for ICSS with the stimulation parameters used in the modified-LI5 task to assess the possible reinforcing effect of the laser stimulation. During 8 1 h daily sessions, rats had access to two nosepoke ports; a response at the active nosepoke resulted in delivery of a 0.5 s train of light pulses (20 Hz, 5 ms pulse duration). Inactive nosepokes were recorded but had no consequence. Eight additional 1 h ICSS sessions were conducted in which a response in the active nosepoke resulted in a 2 s laser stimulation (20 Hz, 5 ms duration) paired with a cue light in the active nosepoke. Correct ICSS in these conditions (above 150 responses at the active nosepoke) was used as criteria for rats' inclusion in the optogenetic experiment, in addition to correct virus expression and optic fiber placement in the VTA.

#### Tissue collection

Rats were deeply anesthetized with sodium pentobarbital and transcardially perfused with cold phosphate buffered (PB) saline (PBS) followed by 4% paraformaldehyde (PFA). Brains were removed and post-fixed overnight in 4% PFA. Brains were then cryoprotected in a PB solution containing 30% sucrose for at least 48 h, frozen into liquid isopentane and store at -80°C until processed for histology.

#### Histology

Brains were processed into 50 µm-thick coronal sections on a cryostat (Leica Microsystems). To verify viral expression of dLight and optic fiber placement, brain slices were directly mounted on microscope slide and coverslipped with Vectashield mounting medium containing DAPI. Brain sections were then imaged with a Zeiss Axio 2 microscope. To verify viral GCaMP, ChR2 and eYFP expression in midbrain dopamine neurons, we performed immunohistochemistry for tyrosine hydroxylase (TH) and GFP (for GCaMP only). Sections were washed in PBS and incubated with bovine serum albumin (BSA) and Triton X-100 (each 0.2%) for 20 min. 10% normal donkey serum (NDS) was added for a 30 min incubation, before primary antibody incubation (mouse anti-GFP, 1:1500, Invitrogen; rabbit anti-TH, 1:500, Fisher Scientific) overnight at 4°C in PBS with BSA and Triton X-100 (each 0.2%). Sections were then washed with PBS and incubated with 2% NDS in PBS for 10 min. Secondary antibodies were added (1:200 Alexa Fluor 488 donkey anti-mouse; 1:200 594 donkey anti-rabbit, Invitrogen) for 2 h at room temperature. Sections were then washed in PBS and in PB and mounted with Vectashield mounting medium containing DAPI. Fluorescence as well as optic fiber tracks were then visualized. In order to determine the specific targeting of TH-Cre$^{+}$ neurons by DIO-viruses, 20 x three-channel images along the medial-lateral and anterior-posterior gradients of the midbrain were taken, using equivalent exposure and threshold settings. With the TH channel turned off, GFP$^{+}$ or eYFP$^{+}$ cells were first identified by a clear ring around DAPI-stained nuclei. The TH channel was then overlaid, and the proportion of GFP$^{+}$ or eYFP$^{+}$ cells co-expressing TH below optic fiber placements was counted.

#### Statistical analysis

##### *Behavioral analysis*

Data were subjected to three-way repeated measures analyses of variance (session as within-factor and task, sex and region as between-subject factors), followed by post-hoc comparisons when indicated, using Student Newman-Keuls pairwise comparison. Significance was assessed against a type I error rate of 0.05. Behavioral data that did not follow a normal distribution were analyzed using non parametric tests (Mann-Whitney Rank Sum t test for group comparison). Statistical tests were conducted on SigmaStat (Systat software Inc., San Jose, USA) and SPSS (IBM, Amorak, NY).

##### *Analysis of photometry recordings*

- ***Signal acquisition***

A real-time signal processor (RP2.1, Tucker-Davis Technologies) running Synapse software was used to modulate the output of each LED and record the photometry signals. The signals were sampled from the photodetector at a rate of 6.1 kHz and demodulated and decimated to 1017 Hz before being saved to disk. For analysis purposes, both signals were downsampled to 102 Hz, and a least-squares linear fit was applied to the 405 nm signal to align it with the 465 nm signal. This fitted 405 nm signal served as the reference for normalizing the 465 nm signal using the formula % ΔF/F =100 * (465 nm signal – fitted 405 nm signal)/(fitted 405 nm signal).

- ***Main analysis across trials during training***

For LI5/LR5 tasks, all ΔF/F photometry traces were analyzed from -2 to 2 s around each event of the behavioral sequence comprising the lever insertion (LI), the sequence of lever presses (LP1 to LP5) and the port entry (PE; reward retrieval). To analyze rapid changes in activity across training sessions, trials from all recording sessions were concatenated and the mean activity across the first 15 trials, the next 15 trials (trials 16-30) and the last 15 trials was compared for each task. Unless otherwise specified, the area under the curve (AUC) was computed from 0 to 1 s post-event. For statistical analysis, two-way repeated measure ANOVAs (trial set as within-subject factor and sex as between-subject factor) were conducted. Deviation from baseline (%dF/F ≠ 0) was determined via bootstrapped confidence intervals (99% or 95% CIs) as described in[66]. A consecutive threshold of 17 consecutive 10 ms time bins was applied for statistical evaluation.

- ***Deconvolution analysis***

For analysis purposes, both signals were downsampled to 25 Hz. In order to isolate response to each behavioral event (lever presses and reward retrieval), kernels were calculated as previously described [67]: time-dependent GCaMP6f signal was modeled as the sum of the response to each behavioral event. Calcium response to each event was computed as the convolution of a time series representing the time of the event (series of 0 s and 1 s, in which 1 s correspond to the event timestamp) and to the kernel corresponding to the response profile to that event. The model can be written as follows, where g(t) is the GCaMP6f signal, where a, b, etc., are example of behavioral event time series, and where $k^a$ and $k^b$ are the kernels for the corresponding event:

$$g(t) = g0 + \sum_{t'=-1s}^{2s} a(t-t')k^a(t') + \sum_{t'=-1s}^{2s} b(t-t')k^b(t') + \cdots + error$$

For each recording session, the coefficients of the kernels were solved using the method of least-squares in Python (LinearRegression from sklearn package). **Suppl. Fig. 4** displays individual rats' kernel across the first and last recording sessions in the LI5 and LR5 tasks.

To identify when the signal significantly deviated from baseline, we used bootstrapping to obtain the 99% confidence interval and applied the consecutive thresholds method [66]. More specifically, the lower and upper bound of the confidence interval must be above or below 0 for a least 5 consecutive 40 ms time bins to detect statistical deviation from baseline.

- ***Analysis of GCaMP6f and dLight signal under omission schedule***

Photometry traces of % ΔF/F were analyzed within a time window of -5 to 5 seconds around two key events: the port entry that followed a reward omission or unexpected reward delivery. We computed the AUC from 0.5 to 1 second after the reward retrieval. To capture the delayed signal depression following the omission of the reward, we also calculated the AUC from 1 to 3 seconds after omission. To identify significant differences between the LI5 and LR5 tasks as shown in **Fig. 5**, we conducted a comparison of peri-event waveforms between the two tasks, using the two-sample t-test and bootstrap technique (bootstrap difference distribution of randomly resampled means) as described in[66]. Deviation from baseline (%dF/F ≠ 0) was determined via bootstrapped confidence intervals (95% CIs) as described in[66]. A consecutive threshold of 17 consecutive 10-ms time bins was applied for statistical evaluation of differences between tasks and from baseline.
